# Preprocessing choices affect RNA velocity results for droplet scRNA-seq data

**DOI:** 10.1101/2020.03.13.990069

**Authors:** Charlotte Soneson, Avi Srivastava, Rob Patro, Michael B Stadler

## Abstract

Experimental single-cell approaches are becoming widely used for many purposes, including investigation of the dynamic behaviour of developing biological systems. Consequently, a large number of computational methods for extracting dynamic information from such data have been developed. One example is RNA velocity analysis, in which spliced and unspliced RNA abundances are jointly modeled in order to infer a ‘direction of change’ and thereby a future state for each cell in the gene expression space.

Naturally, the accuracy and interpretability of the inferred RNA velocities depend crucially on the correctness of the estimated abundances. Here, we systematically compare four widely used quantification tools, in total yielding twelve different quantification approaches, in terms of their estimates of spliced and unspliced RNA abundances in four experimental droplet scRNA-seq data sets. We show that there are substantial differences between the quantifications obtained from different tools, and identify typical genes for which such discrepancies are observed. We further show that these abundance differences propagate to the downstream analysis, and can have a large effect on estimated velocities as well as the biological interpretation.

Our results highlight that abundance quantification is a crucial aspect of the RNA velocity analysis workflow, and that both the definition of the genomic features of interest and the quantification algorithm itself require careful consideration.

## Introduction

Single-cell RNA-seq (scRNA-seq) enables high-throughput profiling of gene expression on a transcriptome-wide scale in individual cells. The increased resolution compared to bulk RNA-seq, where only average expression profiles of populations of cells are obtained, provides vastly improved potential to study a variety of biological questions. One such question concerns the dynamics of biological systems, reflected in, e.g., cellular differentiation and development. While such dynamical processes would ideally be studied via repeated transcriptome-wide expression profiling of the same cells over time, this is not possible with current scRNA-seq protocols. Existing analysis methods for so called *trajectory inference* are instead applied to one or several snapshots of a population of cells, assumed to comprise all stages of the trajectory of interest. Many computational methods for trajectory inference from scRNA-seq have been presented in the literature (reviewed by Saelens et al. (2019)). These methods typically use the similarity of the gene expression profiles between cells to construct a (possibly branching) path through the observed set of cells, representing the trajectory of interest. Projecting the cells onto this path then provides an ordering of the cells by so called pseudotime.

A different approach to the investigation of developmental processes in scRNA-seq data instead exploits the underlying molecular dynamics. The feasibility of such an approach is based on the observation that, with several commonly used library preparation protocols, not only exonic, but also intronic and exon/intron boundary-spanning reads are observed (La Manno et al. 2018), and the insight that considering these in combination with the exonic reads would allow for direct inference of developmental relationships among cells. Similar observations, coupled with a simple differential equation model of transcriptional dynamics, were previously used for investigation of pre-mRNA dynamics and transcriptional and post-transcriptional regulation of gene expression in exon arrays (Zeisel et al. 2011) and bulk RNA-seq (Gaidatzis et al. 2015), as well as estimation of transcription, processing and degradation rates in bulk RNA-seq (Gray et al. 2014). For scRNA-seq, La Manno et al. (2018) used the differential equation model of Zeisel et al. (2011), describing the rate of change of unspliced pre-mRNA as well as spliced mRNA molecules, as a basis for their investigations. They defined the *RNA velocity* for a given gene in a given cell, at a given time point, as the instantaneous rate of change of the spliced mRNA abundance. Combining the RNA velocities with the estimated mRNA abundances enables reconstruction of the state of each cell at a timepoint in the near future. With the increased popularity of RNA velocity applications in scRNA-seq studies, several computational tools have been developed, both for the preprocessing of the reads and for the RNA velocity estimation. The original *velocyto* software (La Manno et al. 2018) estimates velocities under a steady-state assumption, and provides both Python and R implementations. More recently, Bergen et al. (2019) relaxed the steady-state assumption and considered the full dynamical model, thereby enabling application of the RNA velocity framework to a broader set of biological systems and states. The dynamical model is implemented in their *scVelo* Python package, which also includes an efficient implementation of the steady-state model.

The input to both *velocyto* and *scVelo* effectively consists of two gene-by-cell count matrices; one representing mRNA (“spliced”) abundances and one representing pre-mRNA (“unspliced”) abundances. In practice, these two types of abundances are typically represented by reads mapping to the exonic and intronic regions of the genome, respectively. *Velocyto* provides functions for parsing a BAM file of aligned reads (obtained by other tools such as *CellRanger* (Zheng et al. 2017)) and generating these two count matrices. Dedicated end-to-end functionality for estimation of spliced and unspliced abundances from raw scRNA-seq reads are available within the *kallisto*|*bustools* software suite (Melsted, Ntranos, and Pachter 2019; Melsted et al. 2019) and in *STARsolo*, the single-cell mode of the *STAR* aligner (Dobin et al. 2013). Furthermore, assuming a properly specified set of reference sequences, the required counts can also be obtained using other general-purpose tools for quantification of droplet scRNA-seq data, such as *alevin* (Srivastava et al. 2019a). To our knowledge, no critical evaluation of the differences between the count matrices generated by these tools, and the effects on the downstream RNA velocity estimates, has been performed to date. In this study, we therefore used four public experimental droplet scRNA-seq data sets, generated with the 10x Genomics platform, to compare spliced and unspliced abundance estimates obtained by *velocyto, kallisto*|*bustools, STARsolo* and *alevin*. Including alternative index definitions and parameter settings, we analyzed each of the four experimental data sets with a total of up to twelve different quantification approaches (tool/parameter/index combinations). We illustrate that the estimates of spliced and unspliced abundances can be strongly affected by the choice of tool, as well as by the delineation of exonic and intronic regions; in particular, how genomic regions that are exonic in some annotated transcript isoforms and intronic in others are treated. Moreover, we show that abundance estimation is a critical step of the analysis workflow and that differences at this stage can have considerable effects on both the RNA velocity estimates and subsequent biological interpretation.

## Methods

### Data

In this study, we used four public single-cell RNA-seq data sets, generated by different laboratories using popular droplet-based protocols from 10x Genomics. Three of the four data sets (Pancreas, Dentate gyrus and Spermatogenesis) comprise cells from dynamically developing systems, while the fourth (PFC) contains differentiated cells from adult mouse brain and was chosen as a negative control data set, assumed to not harbor a strong dynamic signal.

- The Pancreas data set (Bastidas-Ponce et al. 2019) stems from a study of endocrine development in mouse, and was acquired with the 10x Genomics Chromium platform, using v2 chemistry. We downloaded the FASTQ files containing the reads from the cells at stage E15.5 from the Gene Expression Omnibus, accession number GSE132188. The RNA read length is 151 nt. A subset of this data set was used for illustration by Bergen et al. (2019), and is included as an example data set in the *scVelo* package. After the respective quantifications, we retained only the cells that are also included in the *scVelo* example data set, from which we also retrieved cell type labels. The final processed data set used for our analyses contains 3,696 cells.
- The Dentate gyrus data set (Hochgerner et al. 2018) considers the developing mouse dentate gyrus, and was acquired with 10x Genomics v1 chemistry. The individual FASTQ files for cells from P12 and P35 were downloaded from the Gene Expression Omnibus, accession number GSE95315, and the reads from each time point were combined in a single pair of FASTQ files, with cell barcodes and UMI sequences in one file and the read sequence in the other. The RNA read length is 98 nt. Similarly to the Pancreas data set, the Dentate gyrus data set is also available as an example data set in *scVelo*, and was used for illustration by Bergen et al. (2019). Only cells that were also studied in the *scVelo* paper were retained for our analysis, and cell type labels were obtained from the *scVelo* example data set. *CellRanger* and *velocyto* were not run on this data set since the downloaded FASTQ files were not in the format expected by these tools. The final processed data set used for our analyses contains 2,914 cells.
- The Spermatogenesis data set (Hermann et al. 2018) consists of steady-state spermatogenic cells from an adult mouse, and was processed with 10x Genomics Chromium v2 chemistry. The submitted BAM file was downloaded from the Gene Expression Omnibus, accession number GSE109033 (sample accession number GSM2928341), and converted to FASTQ format using the *bamtofastq* utility (v1.1.2) from 10x Genomics (https://support.10xgenomics.com/docs/bamtofastq). The RNA read length is 100 nt. Cell type labels were obtained from the corresponding loupe browser file provided by the authors, downloaded from https://data.mendeley.com/datasets/kxd5f8vpt4/1. Only cells that were also included in this file were retained for further analysis after quantification. The final processed data set used for our analyses contains 1,829 cells.
- The PFC data set (Bhattacherjee et al. 2019) consists of cells from the prefrontal cortex of an adult mouse, and was generated using 10x Genomics Chromium v2 chemistry. Since only limited dynamics is expected in this data set, it is used here as a negative control. The submitted BAM file was downloaded from the Gene Expression Omnibus, accession number GSE124952 (sample accession GSM3559979), and converted to FASTQ format using the *bamtofastq* utility (v1.1.2) from 10x Genomics (https://support.10xgenomics.com/docs/bamtofastq). The RNA read length is 98 nt. Only cells annotated to ‘PFC sample 2’ were used, and cell type labels assigned by the data generators were obtained from the GEO record metadata. The final processed data set used for our analyses contains 1,267 cells.

### Feature sequence extraction

All analyses are based on reference files from Gencode, mouse release M21 (Frankish et al. 2019). The desired output from each quantification method is a pair of count matrices; one containing ‘spliced’ or ‘exonic’ counts, and the other containing ‘unspliced’ or ‘intronic’ counts for each gene in each cell. For simplicity, in the remainder of this paper, the terms ‘spliced’ and ‘exonic’ will be used interchangeably to refer to the counts representing the processed mRNA abundances, and ‘unspliced’ and ‘intronic’ counts will similarly refer to the counts representing the unprocessed pre-mRNA abundances.

To enable this type of quantification with *alevin* and *kallisto*|*bustools* (as opposed to the more standard, gene-level expression quantification), the Gencode reference files were processed as follows (also summarized in Table 1). First, we used the *BUSpaRse* R/Bioconductor package v1.0.0 (Moses and Pachter 2019) to extract transcript and intron sequences from the genome sequence and the Gencode annotation GTF file. While the definition of the transcripts is unambiguous, the *BUSpaRse* package supports two ways of defining the introns (see Fig. S1 for a schematic). The ‘separate’ approach considers each transcript separately when extracting the intronic regions (and thus, an intron can overlap with an exonic region of an alternate transcript), while the ‘collapse’ approach first collapses the isoforms of a gene (taking the union of all the exonic regions) before defining the introns as any regions of the gene locus that are not exonic in any of the annotated transcript isoforms of the gene. In effect, the ‘separate’ approach thus considers exonic and intronic regions on an equal footing, while the ‘collapse’ approach represents a prior belief that an ambiguous read is more likely to come from an exon than from an intron. A flanking sequence of length *L* − 1 (where *L* is the RNA read length of the respective study) is added on each side of each intron to account for reads mapping across exon/intron boundaries. For comparison, we also reimplemented the extraction of transcript and intron sequences (for both the ‘separate’ and ‘collapse’ approaches) directly using functions from the *GenomicFeatures* and *BSgenome* R/Bioconductor packages (Pagès 2019; Lawrence et al. 2013). Code used to extract the features has been included in the *eisaR* package (https://github.com/fmicompbio/eisaR), from v0.9, for convenience. In each case, the extracted transcript and intron sequences were written to a joint FASTA file, summarized in Table 1. Upon comparison of the two implementations, we noticed that the current release version (v1.0.0) of *BUSpaRse* returned erroneous feature sequences for multi-exonic transcripts on the negative strand (Fig. S2). For this reason, the features extracted by *BUSpaRse* were not used for further analyses.

**Table 1:** Summary of reference FASTA files containing exonic and intronic sequences. Only files generated by *eisaR* were used for creation of quantification indices.

| FASTA file name | Sequence extraction | Description |
| --- | --- | --- |
| BUSpaRse-separate | *BUSpaRse* | Introns extracted separately from each transcript isoform |
| BUSpaRse-collapse | *BUSpaRse* | Introns extracted after collapsing all transcripts of a gene |
| eisaR-separate | *eisaR* | Introns extracted separately from each transcript isoform |
| eisaR-collapse | *eisaR* | Introns extracted after collapsing all transcripts of a gene |

### Reference index generation

The combined transcript and intron FASTA files were used to build the following quantification indices (summarized in Table 2):

**Table 2:** Summary of reference indices. The full index name is constructed by concatenating the index type to the FASTA file name, e.g. salmon-joint-eisaR-separate refers to the salmon-joint index built for the sequences in the eisaR-separate FASTA file.

| Index type | Index building | Input reference files<br>(Table 1) | Target sequences | Decoy<br>sequences |
| --- | --- | --- | --- | --- |
| salmon-joint | *Salmon* | eisaR-separate,<br>eisaR-collapse | spliced transcripts<br>and introns | genome |
| salmon-spliced | *Salmon* | transcriptome | spliced transcripts | genome |
| salmon-spliced-unspliced | *Salmon* | transcriptome +<br>unspliced<br>transcripts | spliced and<br>unspliced<br>transcripts | genome |
| salmon-spliced-decoy | *Salmon* | eisaR-separate,<br>eisaR-collapse | spliced transcripts | introns +<br>genome |
| salmon-introns-decoy | *Salmon* | eisaR-separate,<br>eisaR-collapse | introns | spliced<br>transcripts +<br>genome |
| kallisto-joint | *kallisto* | eisaR-separate,<br>eisaR-collapse | spliced transcripts<br>and introns | N/A |
| cellranger | *CellRanger* | genome and GTF<br>file | whole genome | N/A |
| star | *STAR* | genome and GTF<br>file | whole genome | N/A |

- a joint transcript and intron index for *Salmon* (v1.0.0) (Patro et al. 2017)
- an index for *Salmon*, considering the transcripts as the features of interest and providing the introns as decoy sequences (Srivastava et al. 2019b)
- an index for *Salmon*, considering the introns as the features of interest and providing the transcripts as decoy sequences
- a joint transcript and intron index for *kallisto* (v0.46.0) (Bray et al. 2016)

In addition to the indices based on transcripts and introns, we built one *Salmon* index from the original Gencode FASTA file with the annotated transcripts, and one *Salmon* index from a FASTA file combining the annotated transcripts and fully unspliced versions of all transcripts. For all *Salmon* indices, the complete genome sequence was provided as a decoy sequence (Srivastava et al. 2019b), with the aim to exclude reads coming from intergenic regions of the genome. Across data sets and reference specifications, this excluded between 1 and 2.5% of the reads from the quantification. For the quantification based on spliced-only transcripts, in which the genome decoy would also capture unambiguously intronic reads, this number was between 2 and 10%. Finally, we built an index for *CellRanger* (v3.0.2) (Zheng et al. 2017) and one for *STAR* (v2.7.3a) (Dobin et al. 2013) based on the reference genome and GTF file from Gencode. The splice junction database overhang in the *STAR* index was set to 150nt, which is at least as long as the read length minus one for all data sets considered here.

To investigate the effect of the choice of flank length in the intron definition, we further built *Salmon* indices (using the ‘separate’ intron definition) with flank lengths equal to the read length minus 21bp, and the read length minus 41bp. We also built an index for *kallisto*|*bustools* using the *kb-python* wrapper (Bray et al. 2016; Melsted et al. 2019), which uses the ‘separate’ intron definition and sets the flank length to 30 bp.

### Sequence uniqueness estimation

Finally, to aid in the interpretation, we estimated the sequence uniqueness for each gene, relative to all other genes, separately for each FASTA file generated as described above (Table 1). The gene uniqueness was defined as the fraction of unique k-mers in the gene, that is, the fraction of the constituent k-mers that are not found in any other gene. For each data set, the k-mer length was set to be equal to the RNA read length minus 1. The sequence uniqueness for a gene was calculated in two different ways. In each case, the full FASTA file with transcript and intron sequences was used as input. First, we estimated separate uniqueness values for the exonic and intronic parts of a gene, by assigning the transcripts and introns of a gene to distinct gene IDs in the uniqueness calculations. With this approach, k-mers that are exonic in one transcript and intronic in another (even if these are isoforms of the same gene) are classified as non-unique, and thus the uniqueness score of an exon sharing some of its sequence with an intron of another transcript from the same gene is reduced. Second, we estimated an overall gene uniqueness, by again considering all (exonic and intronic) sequences, but not penalizing shared sequence between introns and exons of the same gene.

## Quantification

### alevin

For each of the *Salmon* indices described above, we ran *alevin* (v1.0.0) (Srivastava et al. 2019a) to estimate exonic and intronic abundances for each annotated gene. It is worth noting that for abundances obtained with the salmon-spliced-unspliced index, ‘exonic’ and ‘intronic’ abundances refer directly to spliced mRNA and unspliced pre-mRNA abundance estimates, respectively, while for all other indices, ‘exonic’ abundances refer to exonic regions and ‘intronic’ abundances to intronic regions. This affects, for example, how reads aligning completely within an exon are used in the quantification: For the salmon-spliced-unspliced index, such a read could stem from either the spliced or the unspliced transcript (since also the latter contains the exons) and could contribute to the abundance of either (or both) of them, while with the other indices, it will be considered ‘exonic’.

For the *alevin* quantifications, the transcripts and introns (or unspliced transcripts) from the same gene were manually annotated with different gene IDs, in order to obtain separate exonic and intronic gene-level abundances despite estimating them jointly. For the indices with decoys, the exonic gene counts were defined as the counts obtained when quantifying against the transcript index (with introns as decoys) and the intronic gene counts were similarly obtained by quantifying against the intron index (with transcripts as decoys). Hence, for these approaches, it is possible for a read that maps equally well to an exonic and an intronic sequence to be included in both the exonic and the intronic count matrices, and thus be counted twice.

### kallisto|bustools

For each of the *kallisto* indices, we applied *kallisto*|*bustools* (v0.46.0) (Melsted, Ntranos, and Pachter 2019; Melsted et al. 2019) to generate a BUS file. Barcodes were corrected using the list of available cell barcodes from 10x Genomics for the appropriate chemistry version, and the BUS file was sorted using *kallisto*|*bustools*. Next, the BUS file was subset with the capture command of *kallisto*|*bustools* to generate separate BUS files to be used for the quantification of exonic and intronic features, respectively. The gene-level exonic and intronic counts were subsequently obtained using the count command. The capture was performed using two different approaches:

- ‘include’, where the features of interest for the quantification at hand are provided to *kallisto*|*bustools*. In other words, the transcript IDs are provided as the -c argument to quantify the exonic abundances, and the intron IDs are provided to quantify the intronic abundances. In practice, this means that reads in equivalence classes containing at least one transcript are retained for the exonic quantification, and reads in equivalence classes containing at least one intron are retained for the intronic quantification. Hence, equivalence classes containing both exonic and intronic features will be provided to both the exonic and intronic quantification steps.
- ‘exclude’, where the features that are not of interest for the quantification at hand are provided to *kallisto*|*bustools*, and subsequently excluded. In other words, intron IDs are provided as the -c argument to quantify the exonic abundances, and the transcript IDs are provided to quantify the intronic abundances, and in addition the -x flag is used to indicate that the provided IDs represent sequences to be excluded. In practice, this means that only reads in equivalence classes that don’t contain *any* introns will be retained for the exonic quantification, and only reads in equivalence classes that don’t contain any transcripts will be retained for the intronic quantification. Hence, equivalence classes containing both exonic and intronic features will be excluded in both steps.

In addition to the manual application of *kallisto*|*bustools* as described above, we applied the *kb-python* wrapper for quantification based on the corresponding index. With its default settings, it calls *kallisto*|*bustools* with the ‘exclude’ capture approach.

### Velocyto and STARsolo

*CellRanger* (v3.0.2) (Zheng et al. 2017) and *velocyto* (v0.17) (La Manno et al. 2018) were run with default settings to generate exonic and intronic counts based on the *CellRanger* index. *STARsolo* (v2.7.3a) (Dobin et al. 2013) was run using the *STAR* index, specifying the SOLOfeatures argument to generate ‘Velocity’ (exonic and intronic), ‘Gene’ (regular exonic gene expression) and ‘GeneFull’ (reads with any overlap with the gene locus) counts. Based on these count matrices, we obtained exonic and intronic count matrices in two different ways. First, we directly used the ‘Velocity’ count matrices as exonic and intronic counts (below referred to as *starsolo*). Second, we used the ‘Gene’ count matrix as the exonic counts, and the difference between the ‘GeneFull’ and ‘Gene’ counts as the intronic counts (below referred to as *starsolo_diff*). For genes where the ‘Gene’ counts were higher than the ‘GeneFull’ counts, the intronic count was set to zero. This can happen, for example, for a gene located in the intron of another gene. In the ‘GeneFull’ quantification, reads mapping to such a gene are considered ambiguous and therefore discarded. However, they may be assigned in the ‘Gene’ quantification, if they are compatible with the annotated gene model. An overview of the evaluated quantification approaches is provided in Table 3.

**Table 3:**
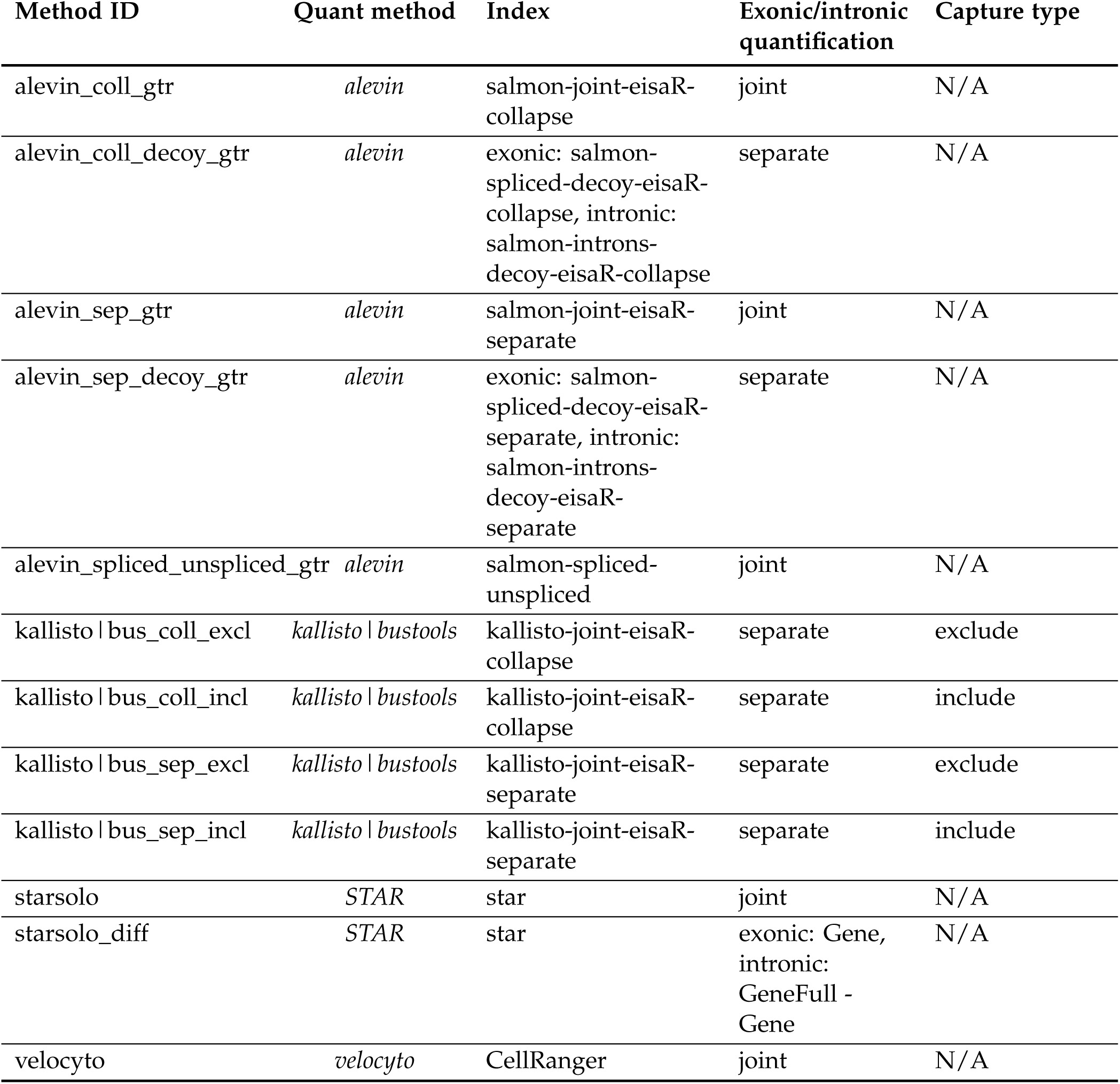
Summary of quantification approaches. In addition to the strategies included in this table, we applied *alevin_sep_gtr* after using different flank length when defining the intronic reference sequences, *alevin_sep_gtr* in unstranded mode, and the *kb-python* wrapper around *kallisto*|*bustools*. The effects of these modifications are evaluated in Fig. S3.

### Cell filtering and data processing

For each quantification, we generated a SingleCellExperiment object (Lun and Risso 2019) containing the exonic and intronic counts. Only cells and genes included by all methods were retained for further analysis. For the Pancreas and Dentate gyrus data sets, we further subset the objects to only the cells analyzed by Bergen et al. (2019), for the PFC data we kept only cells annotated to PFC sample 2, and for the Spermatogenesis data set only cells with an assigned cell type label provided by the data generators were retained. For visualization purposes, we calculated a single low-dimensional representation based on the *alevin* quantification of only the spliced mRNAs (using the original transcriptome FASTA file from Gencode). After normalization with *scater* v1.14.6 (McCarthy et al. 2017), using the library sizes as size factors, we extracted 30 principal components from the log-transformed normalized count values. The *scater* package was then used to apply UMAP (McInnes, Healy, and Melville 2018) with default parameters to the PCA output to obtain a two-dimensional representation that was used for visualization of estimated velocities.

### Visualization

In order to visualize the coverage pattern of reads within genomic regions, we subset the BAM file generated by *CellRanger* to only the reads assigned to the retained cell barcodes, using the *subset_bam* tool v1.0 from 10x Genomics (https://github.com/10XGenomics/subset-bam). Next, we used *BEDTools* v2.27.1 (Quinlan and Hall 2010) to calculate the coverage along the genome, separately for reads on the positive and negative strand. The *bedGraphToBigWig* script from Kent Tools v20190212 (Kent et al. 2010) was used to convert the resulting bedGraph file to bigwig format.

Coverage patterns, together with annotated gene models, were visualized using the *Gviz* R/Bioconductor package v1.30.0 (Hahne and Ivanek 2016). In these figures, the annotated gene models are visualized by their genomic coordinates, together with coverage tracks of reads aligned to the positive and negative strand of the genome. The alignments are aggregated across all the retained cells in the data set. All alignments contained in the BAM file are included; hence, multimapping reads are represented in all the reported mapping locations. Moreover, no UMI deduplication is performed and thus the number of reads reported in the coverage tracks are often higher than the total UMI count returned by any of the counting methods. It is also important to note that while the gene models and coverage tracks are represented with respect to a genomic reference for ease of interpretation, both *alevin* and *kallisto*|*bustools* perform the quantification based on mapping to transcriptomic features, not alignment to the genome. Thus, these plots are not intended to provide an exact correspondence between mapped reads and estimated UMI counts, but rather serve as illustrations to aid in the understanding of the causes of differences between the counts from the various methods.

### RNA velocity estimation

SingleCellExperiment objects with exonic and intronic gene-level UMI counts were converted to Ann-Data objects (Wolf, Angerer, and Theis 2018) using the *anndata2ri* package v1.0 (https://github.com/theislab/anndata2ri). The *scVelo* package v0.1.24 (Bergen et al. 2019) was then used to normalize the counts and select the 2,000 most highly variable genes separately for each quantification approach, after excluding all genes with less than 20 assigned reads across the exonic and intronic components (only summing across cells with nonzero exonic and intronic count). Note that by default, *scVelo* selects highly variable genes based on the spliced counts only. RNA velocity estimates were obtained using the dynamical model implemented in *scVelo*. For comparison, we also performed downstream analysis and visualization of the RNA velocity using only the genes that were selected (and for which *scVelo* returned a finite velocity value) by *scVelo* with all the quantification approaches.

The *scVelo* analysis returns a gene-by-cell matrix of estimated velocities, as well as corresponding matrices of normalized (spliced and unspliced) abundances. Based on these matrices, we estimated both gene- and cell-wise Spearman correlations between the different types of abundances, as well as between the abundances and the velocity estimates. It is worth noting that the velocity calculations are performed separately for each input gene, and the resulting values are therefore independent of which other genes are included in the data set (under the assumption that the normalized abundance values stay unchanged).

Based on the estimated velocity vectors and the differences between the expression profiles of different cells, *scVelo* calculates a cosine correlation (*π*_*ij*_) for each pair of ‘neighboring’ cells. A high value of *π*_*ij*_ indicates that the velocity vector of cell *i* points in the direction from cell *i* to cell *j* in gene expression space. Conversely, if, for a given *i, π*_*ij*_ is small for all *j*, the velocity vector of cell *i* does not point in the direction of any other cell in its neighborhood. With this in mind, we calculate max_*j*_ *π*_*ij*_ for each cell *i* and use this as a proxy for the presence of systematic dynamics in a data set. For each cell, *scVelo* further provides an estimate of the velocity confidence (representing the average correlation of the velocity vector of the cell and those of its neighbors), and an estimated shared (across genes) latent time. The latter was used to contrast the negative control data with the other three data sets, based on the assumption that for a data set without continuous dynamics, the latent time estimates for cells of the same cell type would be more similar to each other than in a data set with a continuous dynamic signal.

### Low-dimensional embedding of velocities

All velocity estimates were embedded into the same low-dimensional representation, calculated from the spliced-only abundance quantification by *alevin* as described above. The embedded velocity vector for cell *i*, calculated by *scVelo*, is given by

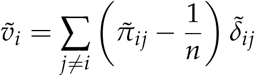

where 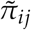 is the transition probability from cell *i* to cell *j* (derived from the cosine correlation *π*_*ij*_), *n* is the number of cells, and

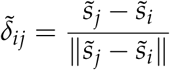

is the normalized difference of the coordinates of cells *i* and *j* in the low-dimensional embedding. The fact that 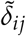 is normalized implies that the length of 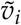 indicates to what extent the cells to which cell *i* has high transition probabilities are all located in the same direction from cell *i* in the low-dimensional representation. It further implies that the embedded velocities are potentially more comparable across methods than the original velocity vectors, since the magnitudes of the latter depend on the normalized abundance levels of the genes, and since the velocity vectors will only be directly comparable between methods if they are based on the same set of input genes. In order to compare the velocity embeddings across methods, we calculate a concordance score for each cell. The score for cell *i* is defined as the ratio between the length of the sum of the embedded velocity vectors for cell *i* across all quantification methods, and the sum of the lengths of the individual embedded velocity vectors. In other words, the score for cell *i* is given by

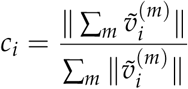

where the sum is taken over all methods *m*, and 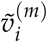 is the embedded velocity vector for cell *i* with method *m*. If all embedded velocity vectors for cell *i* point in the same direction in the low-dimensional representation, this ratio will be close to 1, while if there is less concordance between the different quantification methods, the ratio will be lower than 1.

## Results

### The total UMI count varies between methods

With the aim of characterizing global differences between the counting methods, we first directly compared the total UMI count assigned to exonic and intronic targets by each of the methods. We added up the counts across all cells, either across all genes or within gene subsets stratified by sequence uniqueness (Fig. 1, Pancreas data, similar values were obtained for the other three data sets). There are considerable differences in the assigned UMI counts between the methods. Moreover, these differences are not confined to a small number of susceptible genes, but can be seen across a large fraction of the expressed genes (Fig. S4).

**Figure 1:**
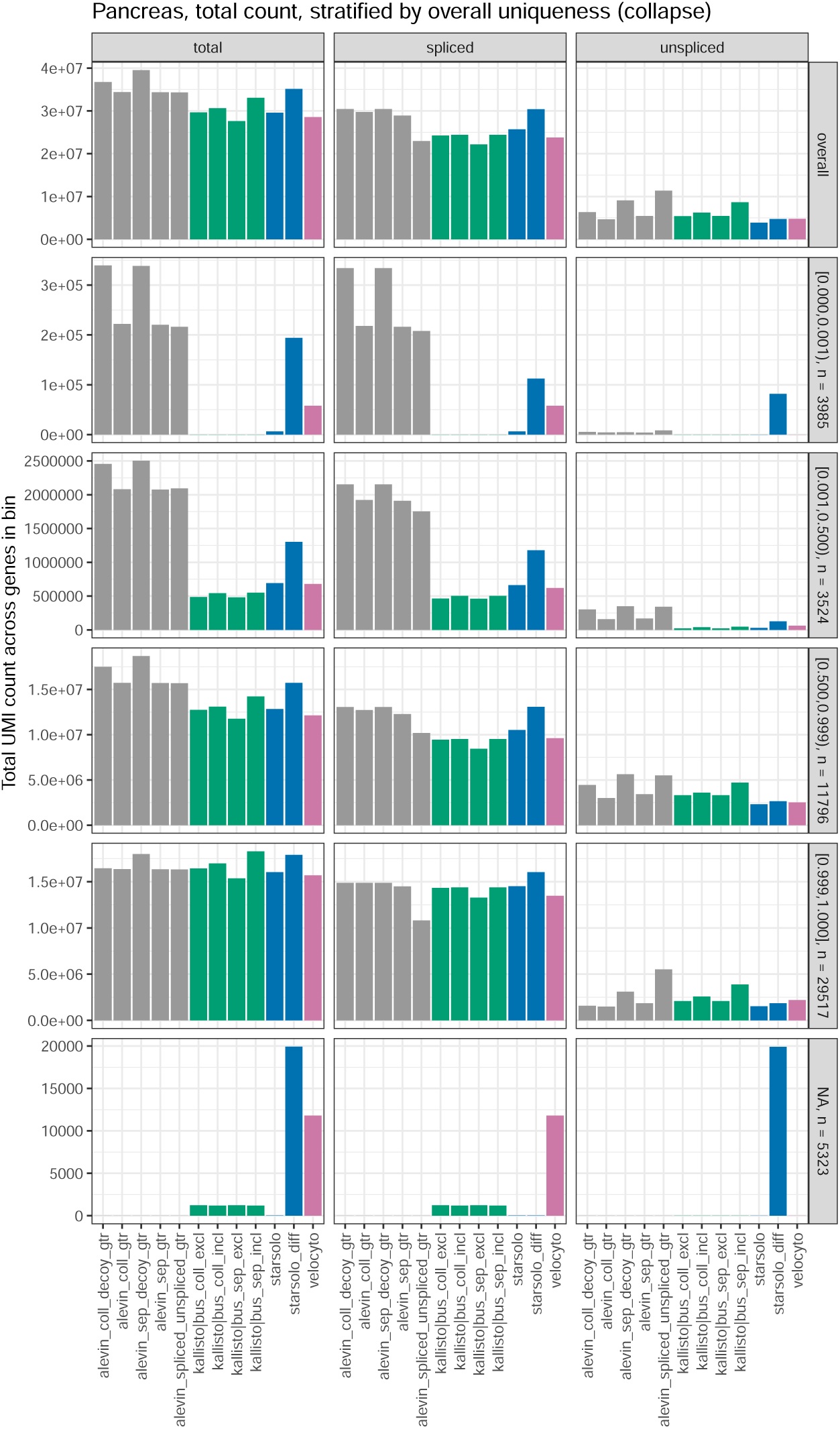
Total UMI count across genes and cells. The bars correspond to the total UMI count, and the split of these into counts for exonic and intronic targets, for each quantification method in the Pancreas data set. Similar patterns were seen in the other three data sets. In addition to the overall count (top row), the figure shows the total count after stratifying genes by the overall fraction of unique k-mers (using the ‘collapse’ annotation), indicated in the vertical panel headers together with the number of genes in the category. The genes for which no uniqueness information could be calculated (the ‘NA’ category) are those for which all transcripts are shorter than the chosen k-mer length (which was set to the read length minus one; here 150nt).

Overall, *starsolo_diff* and the *alevin*-based quantification approaches based on transcript/intron annotations gave the highest total UMI counts, mainly driven by higher counts for the exonic targets. As shown in Fig. 1, this is predominantly due to the assignment of reads to genes with a low fraction of unique k-mers. This is in contrast to *kallisto*|*bustools, velocyto* and *starsolo*, which by default exclude ambiguous reads that map to multiple genes from the quantification.

*Velocyto* and *starsolo_diff* gave the highest UMI counts for genes whose transcripts are all shorter than the read length (genes in the ‘NA’ category in Fig. 1, for which no uniqueness could be calculated since all transcripts were shorter than the employed k-mer length). However, *velocyto* considered most of these reads to be exonic, while *starsolo_diff* assigned them to the intronic features. This behaviour of *starsolo_diff* is likely due to the generation of the intronic counts by subtraction of the exonic count from the full gene locus count. A read which partly overlaps the gene locus but is not consistent with the annotated gene model would be included in the ‘GeneFull’ count but not in the ‘Gene’ count, and thus considered an intronic read, regardless of whether or not the gene actually contains any introns. Similarly, *starsolo_diff* assigned higher counts than both *velocyto* and the default ‘Velocity’ counting of *starsolo*, for both exonic and intronic features.

As expected, the *alevin*-based approaches where exonic and intronic features are quantified separately, as well as the ‘include’ capture mode of the *kallisto*|*bustools* approaches, tend to give higher total UMI count than quantifying exonic and intronic features jointly with *alevin* or running *kallisto*|*bustools* in ‘exclude’ capture mode, especially for the ‘separate’ intron definition. This is likely due to double-counting of some reads that map equally well to an exon and an intron. The difference between the ‘include’ and ‘exclude’ capture approaches is smaller for the ‘collapse’ annotation, since in that case, no genomic regions are annotated as both intronic and exonic for the same gene. The same is true for the difference between the *alevin* quantifications employing joint and separate quantification. It is worth noting that the length of the flanking region chosen when constructing the intronic features for the quantification also influences the counts (Fig. S3). A shorter flank length typically leads to a lower unspliced count, since a larger fraction of the read must overlap the intron for the read to be considered potentially intronic.

In addition to the absolute counts, also the fraction of UMIs assigned to unspliced targets varies between methods, with the largest fraction of intronic counts obtained by *alevin_spliced_unspliced*. This was expected, given that for this method, the ‘intronic’ features are the full pre-mRNA molecules and thus contain both exonic and intronic sequences. Hence, also reads falling in exons may be assigned to the unspliced features.

### Individual genes exemplify methodological differences

Next, we aimed to find individual genes whose count patterns exemplify the main methodological differences among the counting strategies. First, we restricted the set of genes to those that were selected as highly variable by *scVelo*, and thus used for velocity estimation, for at least one counting method. Notably, while a large fraction of these genes were selected across all quantification methods, there were non-negligible differences between the sets of selected genes (Fig. S5). In particular, *alevin_spliced_unspliced* gave the largest number of unique highly variable genes, followed by the *starsolo* variants and *velocyto* depending on the data set. For each of the retained genes, we calculated the fraction of the total counts that were assigned to unspliced features (summarized across all cells). Next, we calculated the standard deviation, across the quantification methods, of these intronic count fractions and selected the top 10% of the genes based on this measure. These genes were partitioned into 10 clusters based on the Pearson correlation dissimilarity (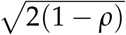 where *ρ* is the Pearson correlation) between the fractions of intron-assigned counts across methods, using hierarchical clustering with complete linkage (Fig. 2). The gene clusters reveal typical cases where the methods yield different exonic and/or intronic counts. Representative genes for each cluster, selected among the genes with the highest correlation with the cluster centroid, are discussed below and illustrated in Figs. S6-S7.

**Figure 2:**
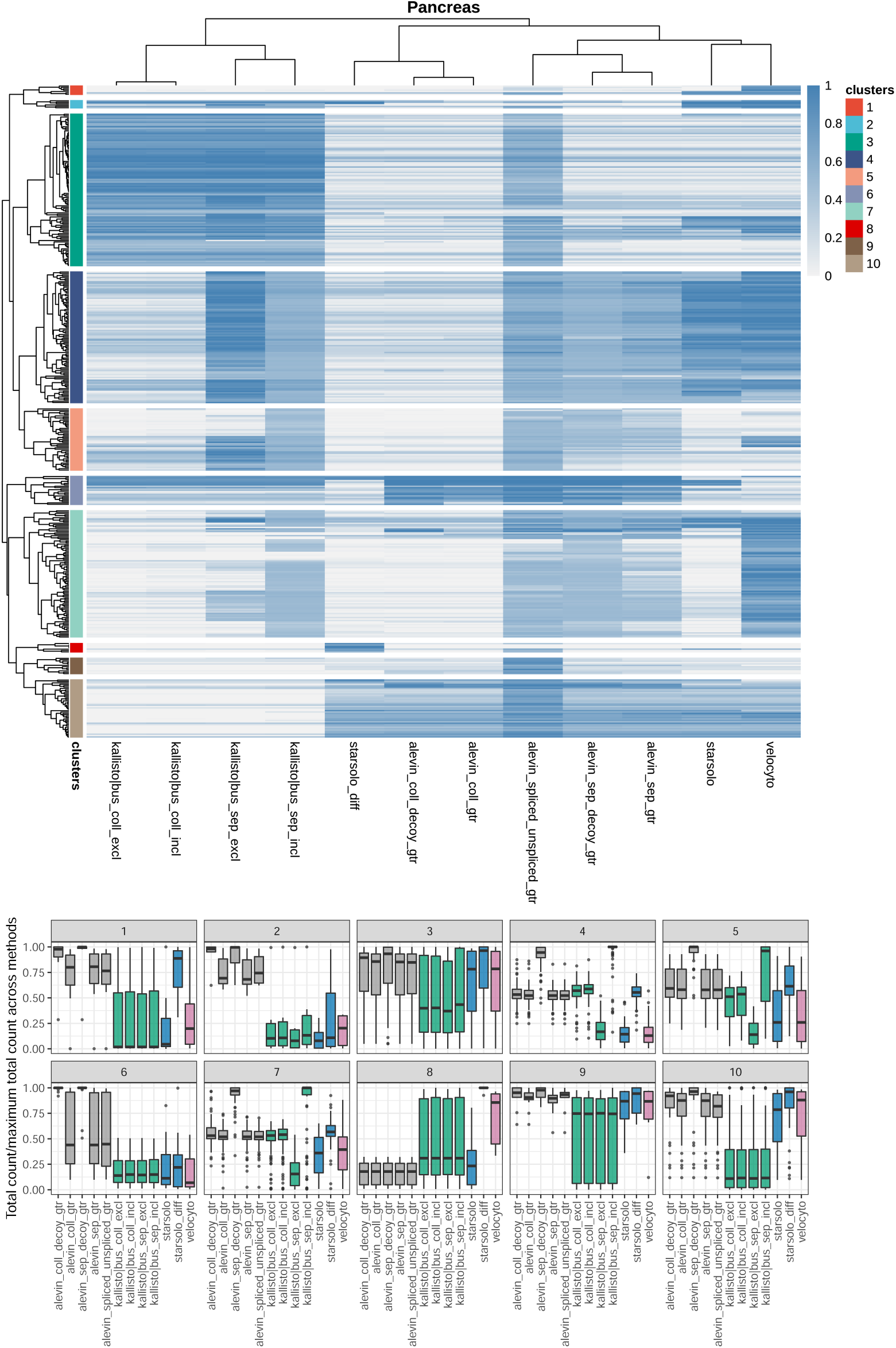
Clustering of genes based on the fraction of unspliced counts. Top panel: Heatmap showing the unspliced UMI count fraction, across all cells, in the Pancreas data. The dendrogram was cut into 10 clusters, indicated by colors to the left of the heatmap. Bottom panel: Relative total count assigned to each gene by the different methods, stratified by gene cluster.

#### Genes with ambiguous regions

(clusters 4, 5, 7, exemplified by Tspan3, Sirt3 and Ssr1 in Fig. S6). For genes in which many of the base positions are annotated to both exons and introns (in different isoforms), the choice of how to define introns (‘separate’ vs ‘collapse’ approaches) has a major effect on the quantifications. If exons are collapsed before the introns are defined, reads falling in ambiguous regions are considered exonic, leading to a higher exonic and a lower intronic count than with the ‘separate’ intron definition. This effect can be seen in approximately half of the genes with the highest variability in the fraction of unspliced counts (Fig. 2), and manifests itself via a low fraction of unspliced reads for the methods based on annotations obtained with the ‘collapse’ intron definition approach. Also *starsolo_diff* falls in this category, since the intronic regions are not considered when the exonic counts are estimated (via the ‘Gene’ count), and thus any read that is compatible with at least one transcript model is considered exonic.

With the ‘separate’ intron definition, running *alevin* with decoys or *kallisto*|*bustools* with the ‘include’ capture double-counts many reads falling completely in ambiguous regions, giving high values of both exonic and intronic counts. Conversely, running *kallisto*|*bustools* with ‘exclude’ capture discards many reads in ambiguous regions, since they will typically be assigned to equivalence classes containing both exonic and intronic targets, and this counting strategy therefore often returns low counts for both types of features. While these effects can be seen in the absolute counts, they do not necessarily affect the ratio of spliced and unspliced counts (Fig. S6).

For genes with many ambiguous regions, *velocyto* often returns a relatively low number of spliced counts, and consequently a large fraction of unspliced counts (clusters 4, 7). This follows from the default ‘permissive’ counting logic where, essentially, a read contributes to the spliced count only if it is consistent with the exonic region of a transcript model, but does not map to an intronic region or an exon/intron boundary of any other transcript model. Also *starsolo* often assigns a low total count for this group of genes.

#### Genes overlapping (introns of) other genes

(clusters 1, 2, 3, 6, 10, exemplified by Chkb, Gm21983, Rassf1, Cnot6 and Tmem120b in Fig. S7). Overlaps between genes can take many different shapes; exons of one gene can overlap either introns or exons of other genes, and the overlap can be on the same or on opposite strands. Reads falling in an exon of one gene and in an intron of another gene on the same strand are considered ambiguous, and are therefore discarded, by *kallisto*|*bustools, velocyto* and *starsolo*. In contrast, *alevin* does not discard the reads, but instead distributes them between the two features. Thus, alevin tends to give a higher fraction of unspliced counts, and in many cases also a higher total count, for genes with other genes in their introns (cluster 6, exemplified by Cnot6 in Fig. S7). It is worth observing that *alevin* will often also assign reads to the gene located within the intron (Gm12191 in Fig. S7, which is the gene in the intron of Cnot6). As for the previous category of genes, *alevin* with the decoy approach double-counts reads mapping equally well to an exon and an intron, regardless of whether or not the exon and the intron belong to the same gene.

In cases of exonic overlap between genes on the same strand (clusters 1 and 2, exemplified by Chkb and Gm21983 in Fig. S7), all methods except *alevin* consider the corresponding reads ambiguous and discards them, leading to a large difference in the total counts (Fig. 2). The main difference between clusters 1 and 2 is that in cluster 1, the gene of interest overlaps partly with an intron of the other gene, and the reads in the overlapping region may be counted by the ‘Gene’ approach of *starsolo-diff*. In cluster 2, the gene of interest overlaps only with exons of the other gene in the region.

Reads falling in overlapping regions of genes *on opposite strands* are not considered ambiguous if the strandedness of the reads is taken into account. However, not accounting for the strandedness (illustrated here by the behaviour of *kallisto*|*bustools*) implies treating such overlaps similar to same-strand overlaps. For example, for a gene located inside the intron of another gene on the opposite strand, all reads not mapping across an exon-exon junction will map equally well to the two genes and thus be discarded (cluster 10, exemplified by Tmem120b in Fig. S7). Exonic overlaps between genes on opposite strands also leads to a decrease in the number of assigned reads if the strandedness is not taken into account (exemplified by Rassf1 in Fig. S7). The assigned count can also increase by performing the quantification in an unstranded manner. For example, reads mapping to the negative strand, not overlapping any feature there but overlapping a gene on the positive strand, can be assigned to the latter. Intronic reads resulting from discordant priming from poly-T sequences, as observed by La Manno et al. (2018), would also be incorporated with an strand-agnostic counting approach. The observation of La Manno et al. (2018) could be reproduced in our data sets (Fig. S8), and the incorporation of opposite-strand reads by *kallisto*|*bustools* is exemplified by Ssr1 and Brsk2 in Fig. S6.

#### Genes with reads only partly overlapping the gene body

(cluster 8, exemplified by 1810019D21Rik in Fig. S6). In some cases, reads extend outside the annotated gene body. The *starsolo_diff* ‘GeneFull’ count incorporates these reads, while the ‘Gene’ count from the same method does not, since they are not compatible with the annotated gene model. As an effect, the difference between them (which is used as the intronic count) can be high, even in cases where the gene does not contain introns, and the total count assigned by *starsolo_diff* is higher than most of the other methods. *Velocyto* and, for some genes in the cluster, the *kallisto*|*bustools* methods, yield a high spliced count, while the *alevin* methods and *starsolo* return lower values.

#### Reads falling in purely exonic regions

Across the clusters, *alevin_spliced_unspliced* often shows a different read assignment compared to the other methods. As previously noted, there is a fundamental difference between the reference used for *alevin_spliced_unspliced* (which considers full spliced and un-spliced transcripts) and those used for the other methods (which consider transcripts and introns). This implies that reads falling completely in exonic regions, from the point of view of *alevin_spliced_unspliced*, are equally likely to come from a spliced as an unspliced molecule, while the other methods consider such reads unambiguously exonic. In practice, this can lead to a lower exonic count and a more even exonic/intronic count ratio for *alevin_spliced_unspliced* than for the other methods. This is exemplified by Map1b (cluster 9) in Fig. S6.

### Large differences in inferred velocities between quantification methods

In the previous sections we showed that there are noticeable differences between the quantification methods, in terms of the total number of UMI counts as well as the distribution of these between spliced and unspliced targets. Next, we asked whether these differences could be seen also in the velocity estimates from *scVelo*, and in the embedding of these in a low-dimensional representation of the cells, which is arguably the most widely used way of interpreting RNA velocity estimates. For our analyses, we provided *scVelo* with raw spliced and unspliced UMI count matrices. These were then filtered and normalized by *scVelo*, and the RNA velocity was estimated for each input gene and each cell. Velocities were estimated for either the individual sets of 2,000 highly variable genes from each quantification method, or the set of genes that were selected by *scVelo* (and obtained a valid velocity value) with all the quantifications.

Interestingly, the estimated velocities consistently showed a lower correlation between methods than the normalized (spliced, unspliced or aggregated) abundances, when calculated across either cells or genes (Fig. S9). Within a cell, there was also a relatively strong correlation between the total gene abundance and the absolute value of the velocity (Fig. S10). This should be factored in when comparing absolute velocities across genes, and it may also suggest that velocity estimates are not directly comparable across quantification methods if the number of assigned reads are very different. For a given gene, the fraction of unspliced counts was also moderately positively correlated with the estimated velocity. The spliced and unspliced abundances were positively correlated for all quantification methods, suggesting that the intronic signal is indeed real and of potential biological relevance, rather than just the result of, e.g., contamination by genomic DNA. Finally, we noticed a moderate positive correlation between the abundance of a gene and the likelihood of the velocity fit in three of the four data sets (exemplified by the Pancreas data set in Fig. S11), while it was substantially lower in the Dentate gyrus data set (Fig. S12).

The velocity estimates were visualized by embedding them into a UMAP representation based on the *alevin_spliced* quantification (note that the *alevin_spliced* counts are not used to estimate the velocity, since no intronic counts are estimated). The UMAP embedding was compared to other types of embeddings (PCA, tSNE, UMAP based on aggregated abundances, unspliced abundances only or spliced and unspliced abundances concatenated), in terms of the length of the embedded velocity vectors as well as the average distance between the velocity vector of each cell and its 10 nearest neighbors. These comparisons suggested that the differences between embeddings were relatively minor, but that UMAP often provided a slightly more interpretable representation (Figs. S13-S14). Embeddings based solely on unspliced abundances were least interpretable from a velocity perspective.

### Differences in velocity estimates directly affect biological interpretation

From the UMAP visualizations it is immediately apparent that the differences in the estimated abundances between the quantification methods directly influence interpretation, e.g., indicated by streamlines pointing in different directions in certain regions of the plots (Figs. S15-S17). These differences are not captured by the velocity confidence estimates returned by *scVelo* for an individual method, which are often high for all the cells (Figs. S18-S20), suggesting that the differences between methods are systematic rather than just the result of random fluctuations or uncertainty in the velocity estimation. The similarity among the low-dimensional velocity embeddings based on different quantification methods increased somewhat when they were derived from the set of shared genes (Fig. 3). However, considerable differences were still seen, indicating that the quantification does not only influence the velocity interpretation via the selection of genes.

**Figure 3:**
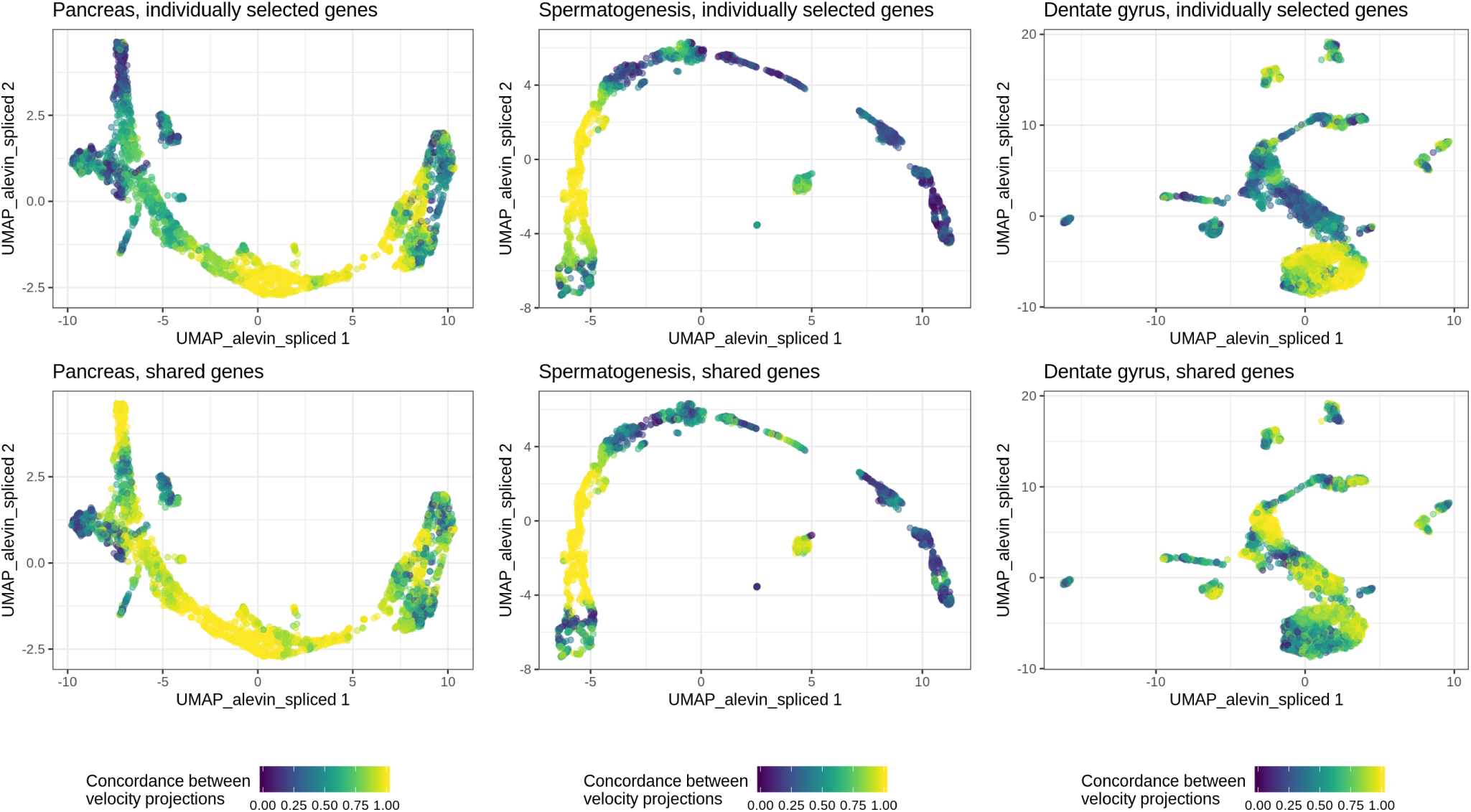
Concordance of velocity projections. Each panel colors the UMAP embedding of the cells by the concordance of velocity projections across methods for one of the three data sets expected to contain systematic dynamics. The concordance for a given cell was defined as the ratio between the length of the sum of the velocity embeddings for the different quantification methods, and the sum of the lengths of the same embedding vectors. A value close to 1 indicates that the embedded velocity vectors all point in the same direction, while a low value indicates larger deviations among them. For each data set, the top panel shows the concordance between the velocity embeddings derived based on all the 2,000 genes selected by *scVelo* for each method, while the bottom panel shows the concordance between the embeddings obtained when only the genes selected for all methods were considered (see also Fig. S5).

The lack of unambiguous ground truth complicates a direct evaluation of the accuracy of velocity estimates from the different quantification methods. In addition, the typical way of interpreting velocity estimates by means of embedded stream lines in a low-dimensional space provides a relatively coarsegrained measure. Nevertheless, Figs. 3 and S15 suggest that for the Pancreas data, the largest differences between methods appear in the differentiated Alpha, Beta, Delta and Epsilon cell types (top left). Here, *alevin_sep_decoy_gtr* and the *kallisto*|*bus* methods induce (partly or fully) a ‘back-flow’, with streamline arrows pointing from the differentiated cells back towards the pre-endocrine cells. A similar observation can be made for *alevin_spliced_unspliced_gtr* and *kallisto*|*bus_sep_incl* for the pre-endocrine cells. The cycling nature of the ductal cells is visible in the embeddings of velocities from most quantification methods, with the exception of *alevin_spliced_unspliced_gtr, alevin_sep_decoy_gtr, kallisto*|*bus_coll_incl*, and *kallisto*|*bus_sep_incl*.

Also for the spermatogenesis data (Figs. 3, S16), the largest differences between methods are seen towards the end of the developmental trajectory. Again, many methods (with the exception of *alevin_sep_gtr* and *kallisto*|*bus_sep_excl*) induce a back-flow, with streamline arrows pointing from the late round spermatids towards the mid round spermatid cluster. In most cases, this back-flow continues through (part of) the mid round spermatid cluster as well.

In the dentate gyrus data, the lowest concordance between velocities based on different quantifications is seen for the cells in the granule cell lineage (middle part). While some quantifications indicate a direction largely from neuroblasts to granule cells (e.g., *alevin_sep_gtr, kallisto*|*bus_coll_incl*), others indicate a strong movement in the opposite direction (e.g., *alevin_sep_decoy_gtr, kallisto*|*bus_sep_incl*). All methods except *kallisto*|*bus_coll_excl, kallisto*|*bus_sep_excl* and *kallisto*|*bus_coll_incl* show a strong dynamic flow within the mature granule cells, and there is further disagreement within the astrocyte cell cluster. Overall, the results from these three data sets highlight that the biological interpretation can be strongly affected by the choice of quantification method.

### Negative control data

The PFC data set was used to compare the methods in terms of their performance on a ‘negative control’ data set, that is, a data set where no strong systematic dynamics are expected. Here, we chose to compare the methods in terms of the maximal cosine correlation between the estimated velocity vector for a cell and the displacement vector to other nearby cells, as calculated by *scVelo*. A low value of this quantity indicates that the velocity vector of a cell does not point in a direction compatible with the difference to any neighboring cell in the data set, which is used here as a proxy for a lack of systematic dynamics. While the maximal cosine correlation varied considerably among cell types (Fig. S21-S24), the value was typically slightly lower for the PFC data set than for the data sets with known dynamics (Fig. 4). The exceptions were *velocyto, alevin_sep_decoy_gtr* and *kallisto*|*bus_sep_incl*, for which the estimated velocities often correlated strongly with the displacement vector to at least one other cell in the negative control data set. We also estimated the standard deviation of the estimated latent times, within each cell type (Fig. S25). In a data set with no continuous trajectories, we would expect a low variation within a cell type (even if there are large differences between cell types). For all methods, the PFC data set indeed showed the lowest variation, as expected.

**Figure 4:**
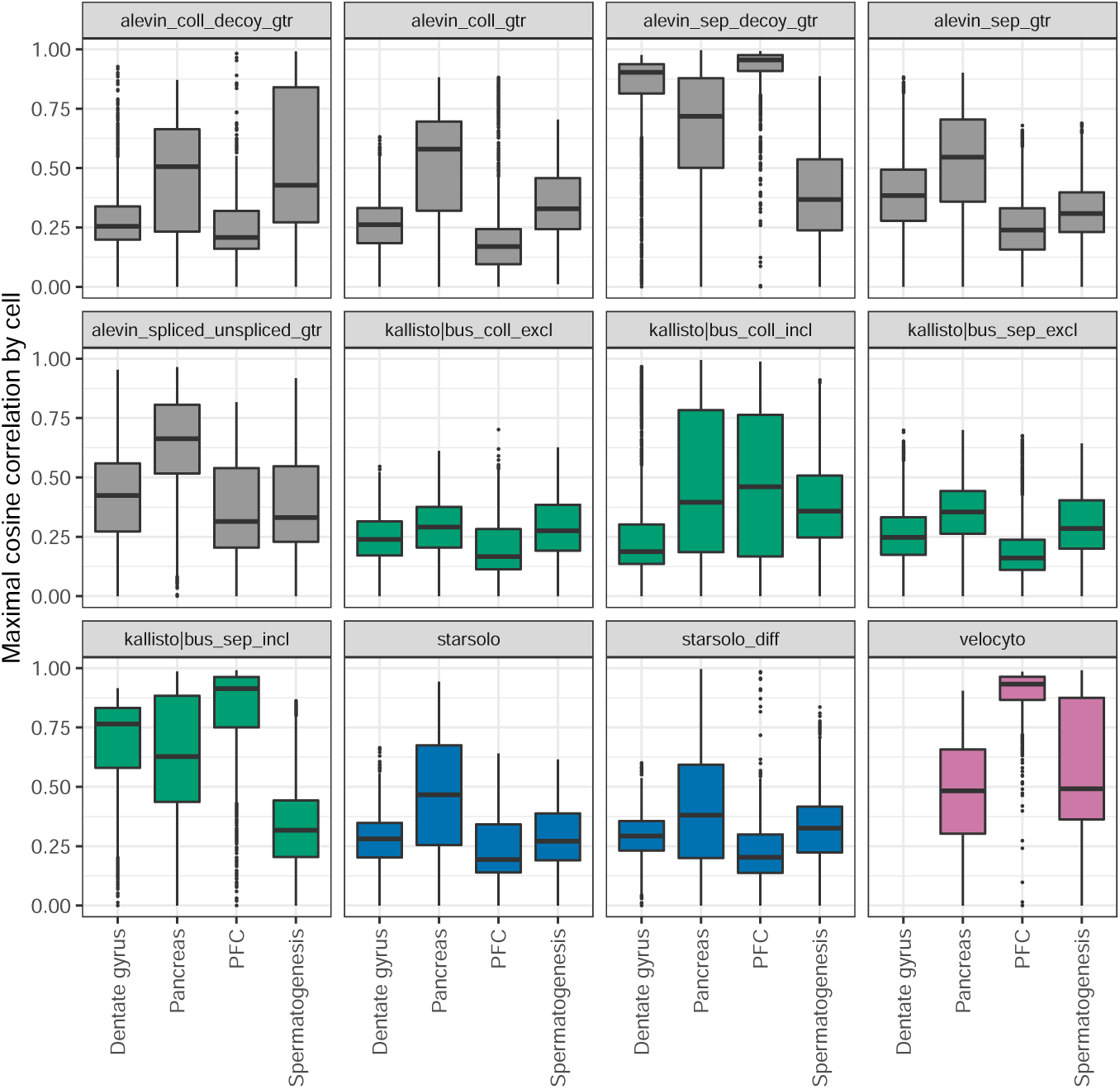
Maximal cosine correlation. The box plots show the distribution, across cells, of the maximal cosine correlation between the velocity vector and the displacement vector to other cells in the neighborhood, estimated by *scVelo* for different quantification methods in the four data sets.

### Discussion and Conclusions

In this study, we have compared different counting strategies for obtaining the spliced and unspliced count matrices required for RNA velocity analysis. Using four experimental droplet scRNA-seq data sets, we have shown that there are considerable differences between the count matrices obtained by different methods that are widely used in the field, and that these differences directly influence the downstream analysis and interpretation of the estimated velocities. This effect is mediated partly by anb impact on the genes that are selected for inclusion by *scVelo*, but differences affecting the interpretation remain even when the same set of genes is used across all methods.

Given the relative immaturity of the RNA velocity field, and the lack of a generally accepted method for generating realistic, simulated data with known ground truth for this application, it is challenging to rank the quantification methods in terms of absolute performance. However, some clear themes emerge from our analysis. First, counting exonic and intronic reads separately, without consideration of whether the read could have resulted from the other type of feature (exemplified here mainly by *alevin_sep_decoy_gtr* and *kallisto*|*bus_sep_incl*) leads to double-counting of reads, and velocities that agree less well with expectations. Second, not considering the strandedness of the reads from 10x Genomics (here exemplified by the *kallisto*|*bustools* variants and by explicitly running *alevin* in unstranded mode) implies that many reads in regions where genes on different strands overlap each other (exonically or intronically) are considered ambiguous. Depending on the method, these reads may consequently be excluded from the quantifications. However, at the same time it provides to ability to include reads resulting from discordant internal priming. Third, deriving the intronic reads by subtracting the ‘Gene’ count from *STARsolo* from the corresponding ‘GeneFull’ count (here denoted *starsolo_diff*) sometimes has unexpected consequences, since the ‘GeneFull’ counting considers all reads that overlap the gene locus, while the ‘Gene’ counting requires that the reads are consistent with the transcript model. Thus, a gene can obtain a nonzero ‘intronic’ count despite completely lacking annotated introns. Moreover, genes located within introns of other genes will often obtain a nominally negative intronic UMI count since non-junction-spanning reads mapping to the former will be considered ambiguous and discarded in the ‘GeneFull’ counting. Fourth, for 3’ tag data such as the 10x Genomics data we have considered in this study, quantifying the spliced and unspliced transcripts (rather than spliced transcripts and introns only) implies that for a large fraction of the reads, it is difficult to resolve whether they stem from the spliced or unspliced target. Thus, this type of reference may be more suitable for full-length scRNA-seq protocols, where reads are sampled across the entire length of the transcript.

Among the counting strategies contrasted in this manuscript, *alevin_sep_gtr, kallisto*|*bus_sep_excl, star-solo* and *alevin_coll_gtr* provided velocity embeddings most in line with expectations in the three real data sets. However, even among these methods, there are large differences in the assigned counts as well as in the handling of ambiguous reads and genomic regions. Going forward, we expect that improvements in counting strategies for scRNA-seq data, specifically tailored for RNA velocity preprocessing, will likely come alongside an increased understanding of the read generation process and the biases underlying specific scRNA-seq library preparation protocols, and that different counting strategies may be optimal for different types of scRNA-seq data. An increased understanding of the read generation process will also enable realistic simulation of sets of spliced and unspliced scRNA-seq reads, which in turn will provide an improved platform for objective evaluation of the performance of counting strategies.

## Data and code access

The code used for our analyses is available on https://github.com/csoneson/rna_velocity_quant. The spermatogenesis data can be downloaded from GEO, accession number GSE109033. The pancreas data set was downloaded from GEO, accession number GSE132188. The dentate gyrus data set was downloaded from GEO, accession number GSE95315. The PFC data set is accessible from GEO under accession number GSE124952 (sample accession number GSM3559979).

## Acknowledgements

The authors would like to thank Dania Machlab, Panagiotis Papasaikas and Michael I Love for helpful discussions and comments on the manuscript.

