## Supplementary Figures for "Preprocessing choices affect RNA velocity results for droplet scRNA-seq data"

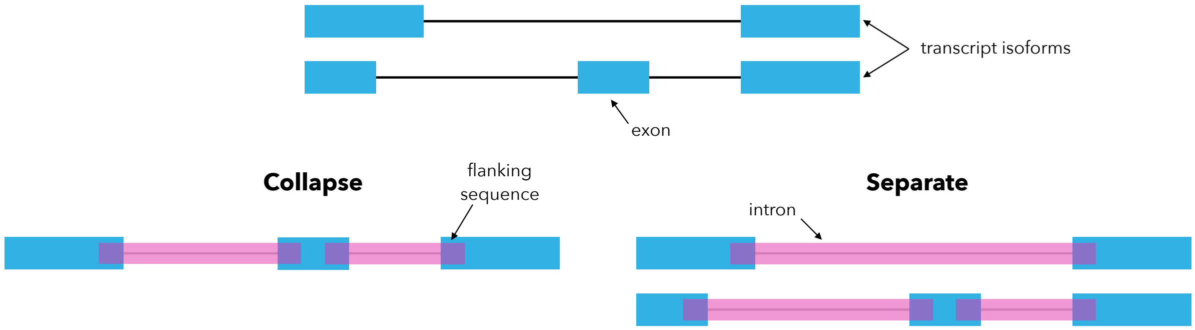


Figure S1: Schematic of the intron definition with the 'collapse' and 'separate' approaches. With the 'collapse' approach, the annotated isoforms of a gene are first collapsed before the introns are defined as any non-exonic region of the gene locus. With the 'separate' approach, introns are separately defined for each isoform. After extracting the intronic regions, a flanking region is added to each side of the introns to accommodate reads overlapping exon/intron boundaries.

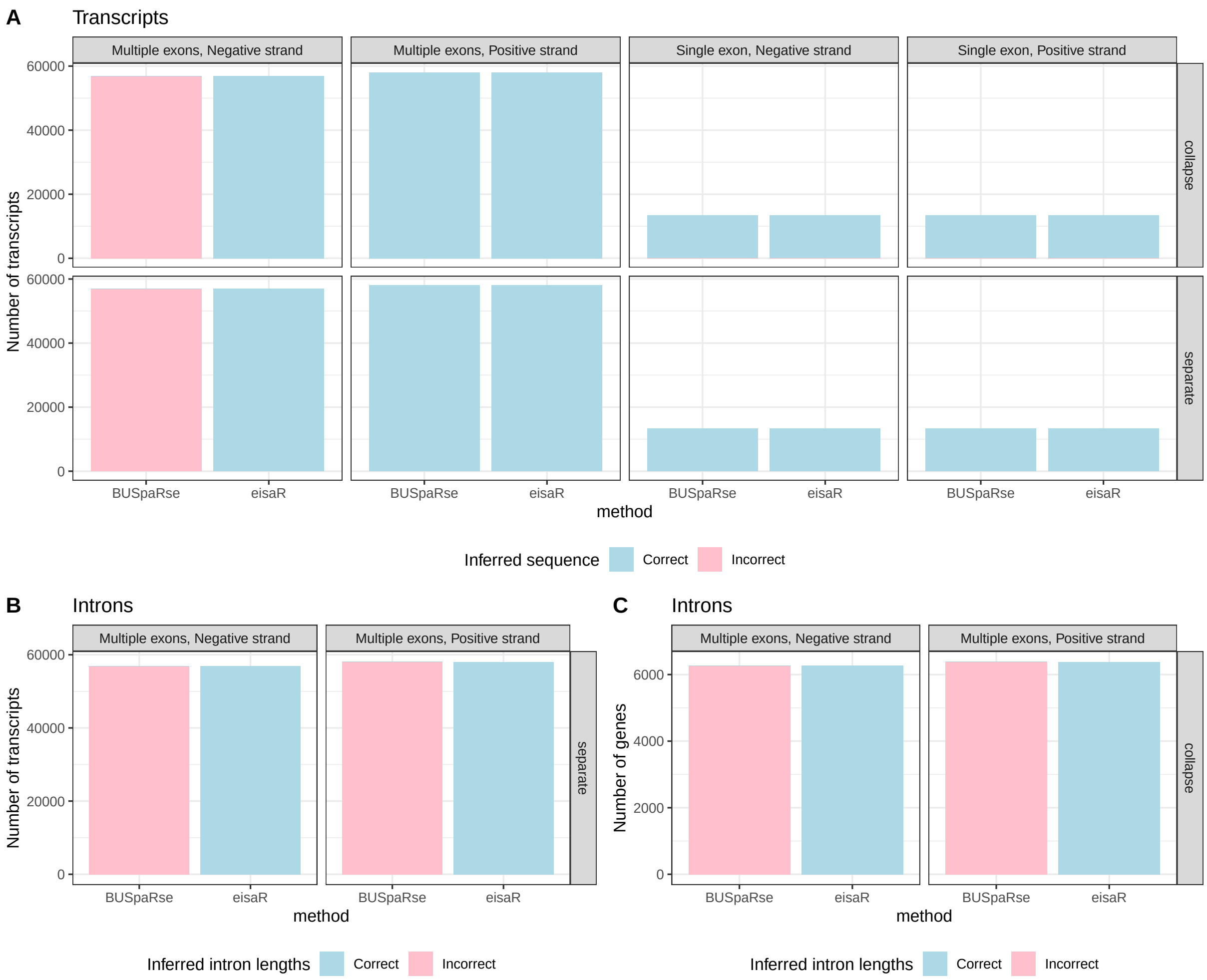


Figure S2: **A**. The number of transcripts extracted by *BUSpaRse* and *eisaR* that have the correct or incorrect sequence, respectively, as determined by comparing to the transcriptome fasta file downloaded from Gencode. Transcripts are stratified by whether or not they are multi-exonic, and by the strand. **B**. The number of transcripts for which the lengths of the introns inferred by *BUSpaRse* and *eisaR* are correct or incorrect, respectively, as determined by comparing to the introns extracted by the `intronsByTranscript` function from the *GenomicFeatures* Bioconductor package ('separate' intron definition). **C**. The number of single-exon genes for which the lengths of the introns inferred by *BUSpaRse* and *eisaR* are correct or incorrect, respectively, as determined by comparing to the introns extracted by the `intronsByTranscript` function from the *GenomicFeatures* Bioconductor package ('collapse' intron definition).

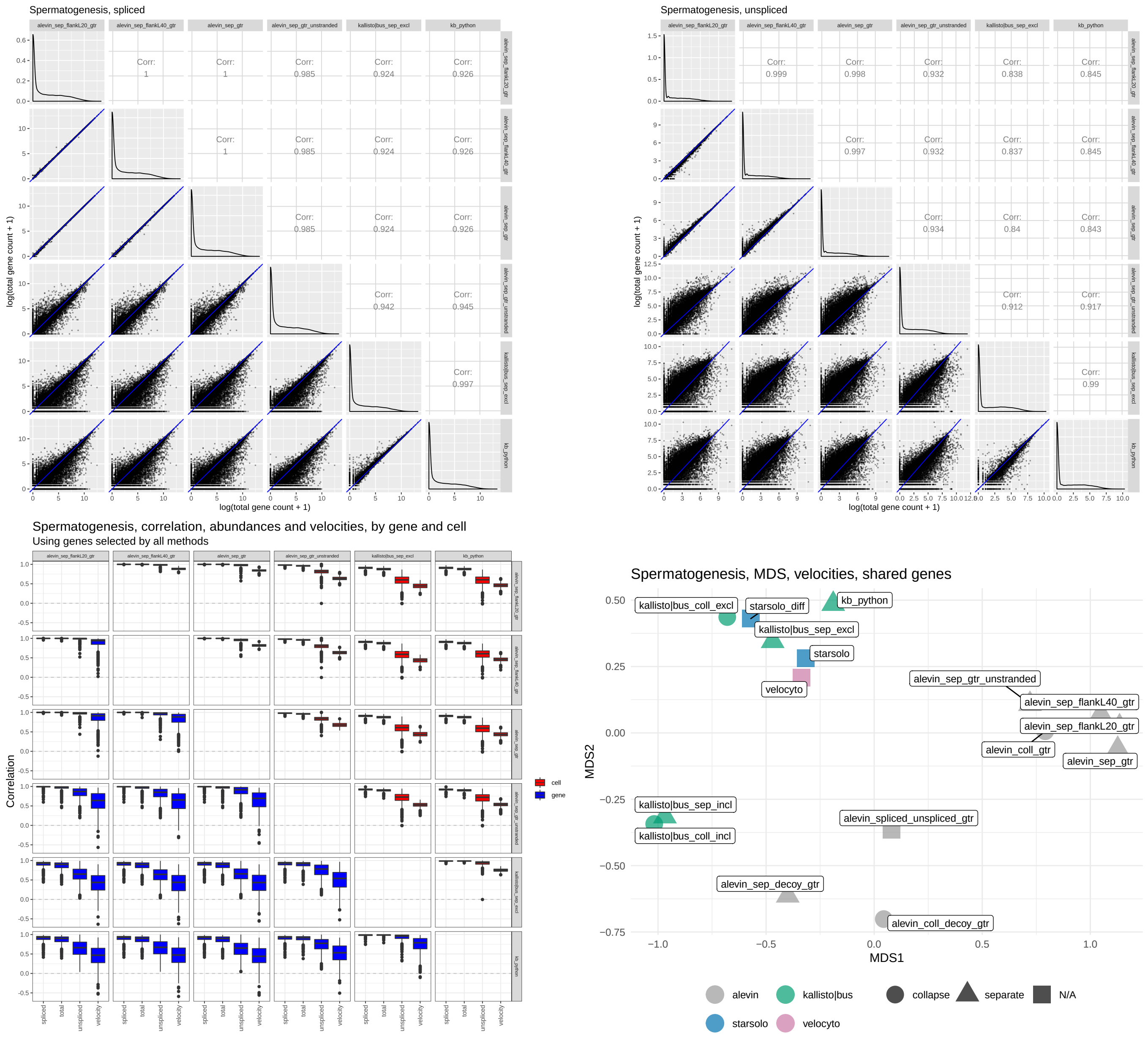


Figure S3: Evaluation of the impact of changing the flank length in the intron extraction, as well as running *alevin_sep_gtr* in unstranded mode, in the Spermatogenesis data set. In each case, the methods are compared to the methodologically most similar among the methods discussed in the main text. *alevin_sep_flankLXX_gtr*, with XX set to either 20 or 40, corresponds to running *alevin* with introns defined using the 'separate' approach, and using a flank length equal to the read length minus (XX+1). The *alevin_sep_gtr* method corresponds to setting XX=0. Further, *alevin_sep_gtr_unstranded* corresponds to running *alevin* in unstranded mode. The *kb-python* wrapper uses the same type of intron definition and capture approach as *kallisto|bus_sep_excl*, but fixes the flank length to 30bp, whereas for *kallisto|bus_sep_excl*, the read length minus 1 was used. Top row: scatter plot of the total spliced and unspliced count assigned to genes with the different methods. Bottom row, left: Spearman correlation between abundances and velocities for each pair of methods. Bottom row, right: A classical multidimensional scaling (MDS) plot based on the Euclidean distances among velocity values for the set of shared genes. The various modifications to the methods have an impact on the derived velocities; however, the modified methods still cluster close to the corresponding base method.

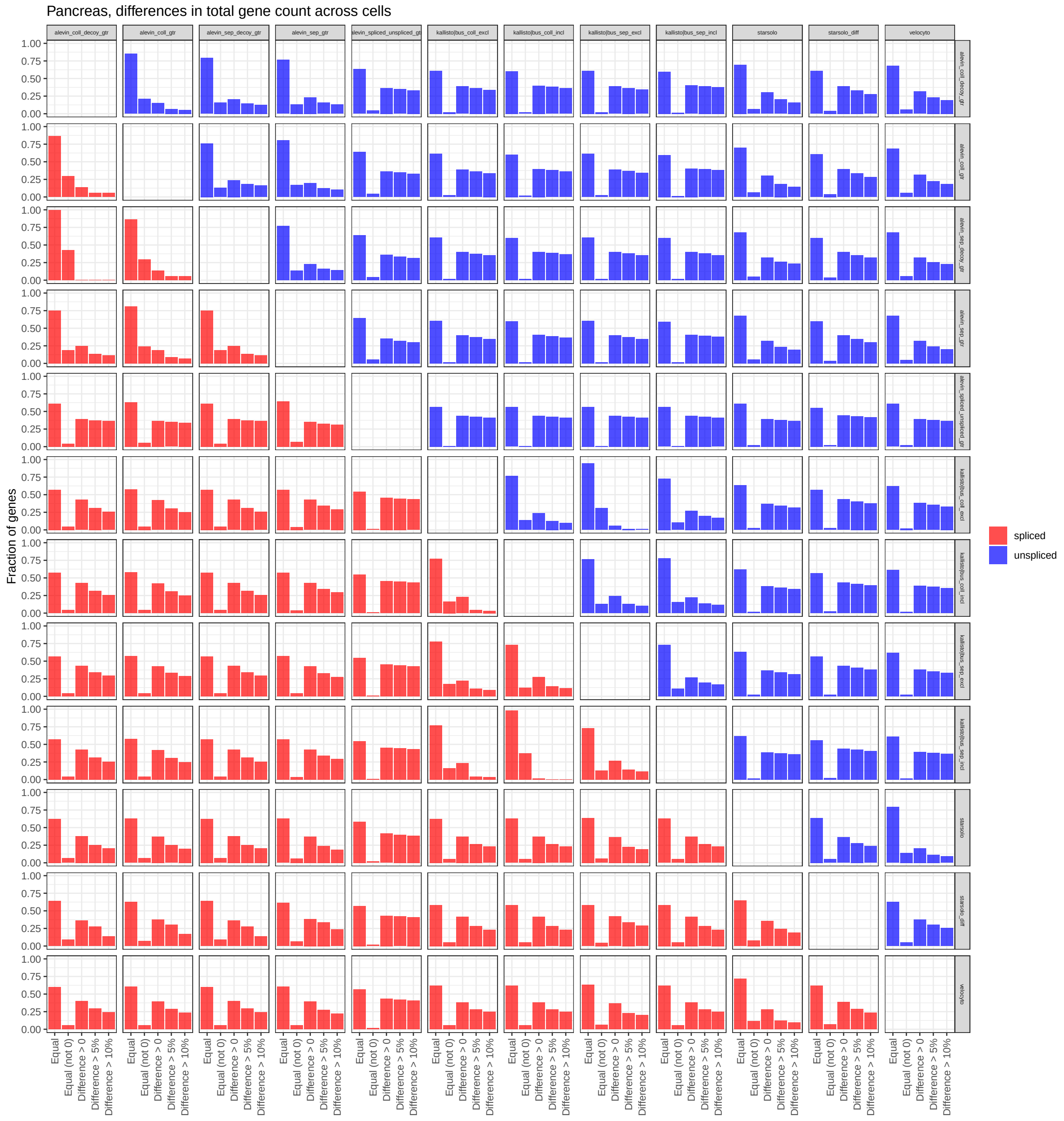


Figure S4: Stratification of genes based on whether or not they obtain the same (spliced/unspliced, respectively) count with different counting methods. For each pair of methods, the figure shows the fraction of genes that obtain the same count with the two methods (both overall and after excluding the genes assigned a count of 0 with both methods) as well as the fraction of genes where the difference between the assigned counts is non-zero, greater than 5% (of the average count assigned by the two methods) or greater than 10%.

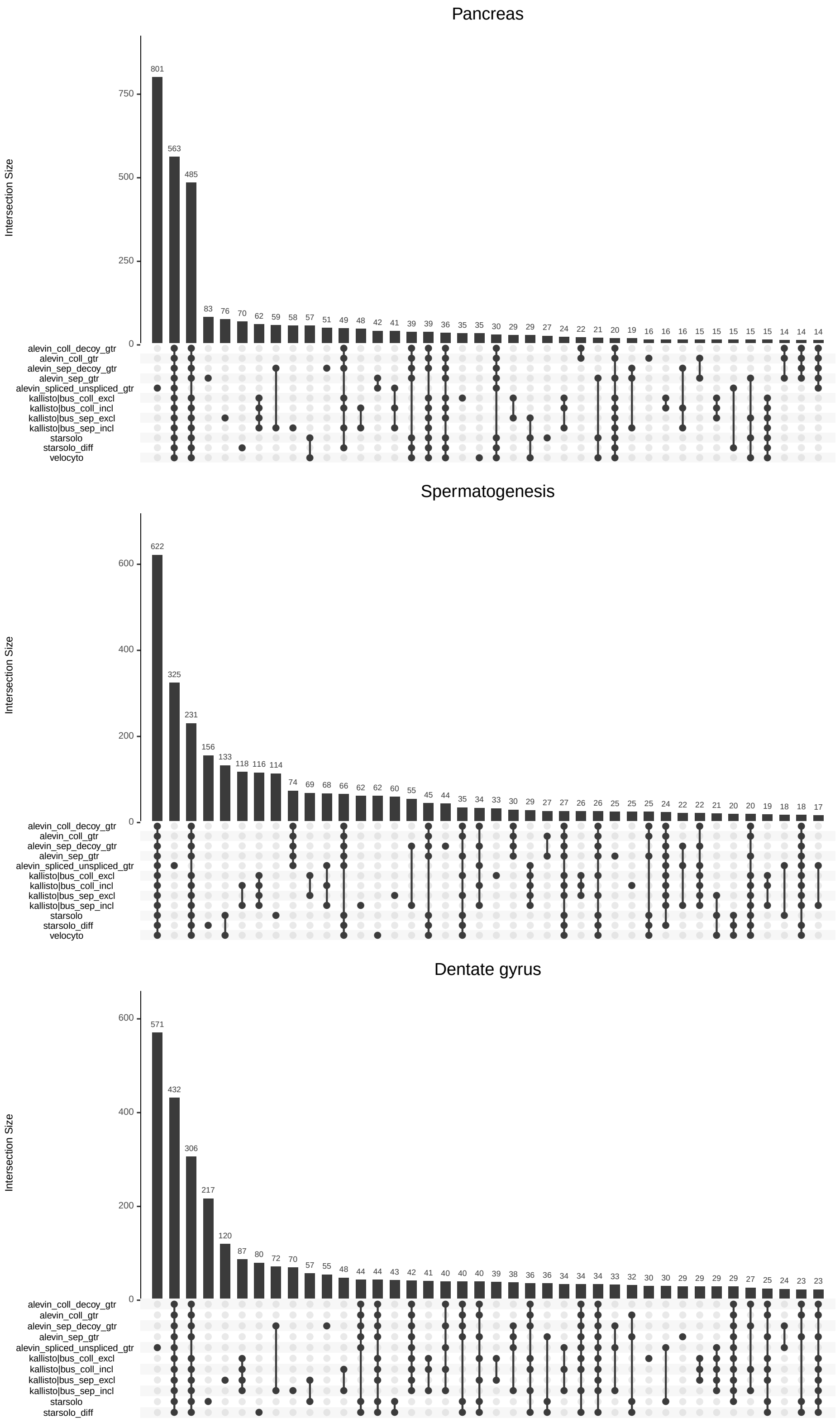


Figure S5: Overlaps among the sets of highly variable genes selected by *scVelo* based on the different quantification approaches. Each column corresponds to the number of genes shared by a particular set of methods (indicated by black dots).

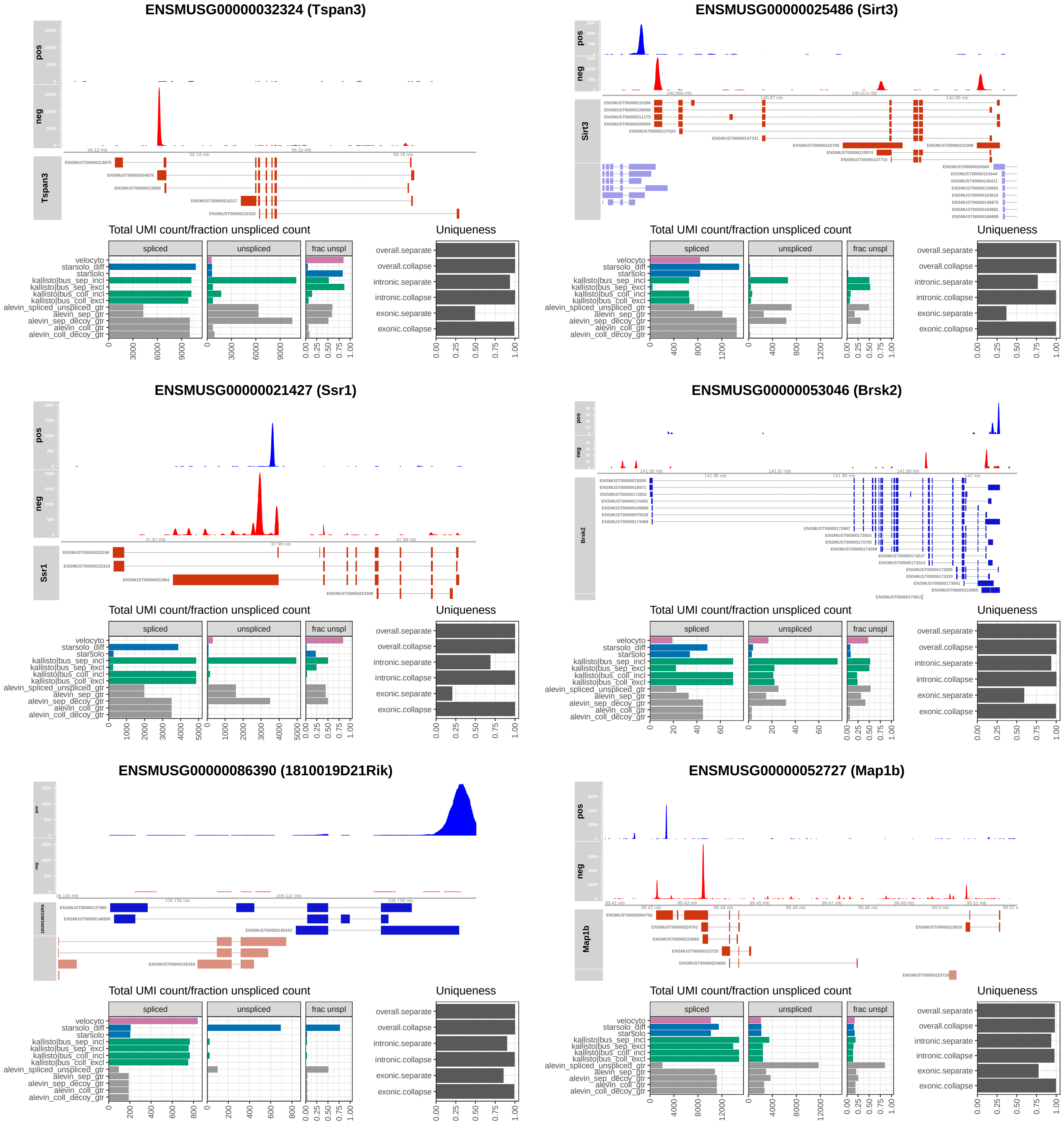


Figure S6: Examples of genes illustrating major differences among the quantification methods. The coverage tracks show the number of reads overlapping each base position, on the positive (blue) or negative (red) strand. The isoforms of the main gene in focus of each panel are similarly shown in blue (positive strand) or red (negative strand). Any overlapping features from other genes are shown in the bottom annotation track, colored in muted blue or red, depending on the annotated strand. The bottom panels show the total exonic and intronic UMI count assigned to the displayed gene by the different quantification methods, as well as the fraction of unspliced counts, and the fraction of unique k-mers in the gene overall, as well as in the exons and introns (see Methods).

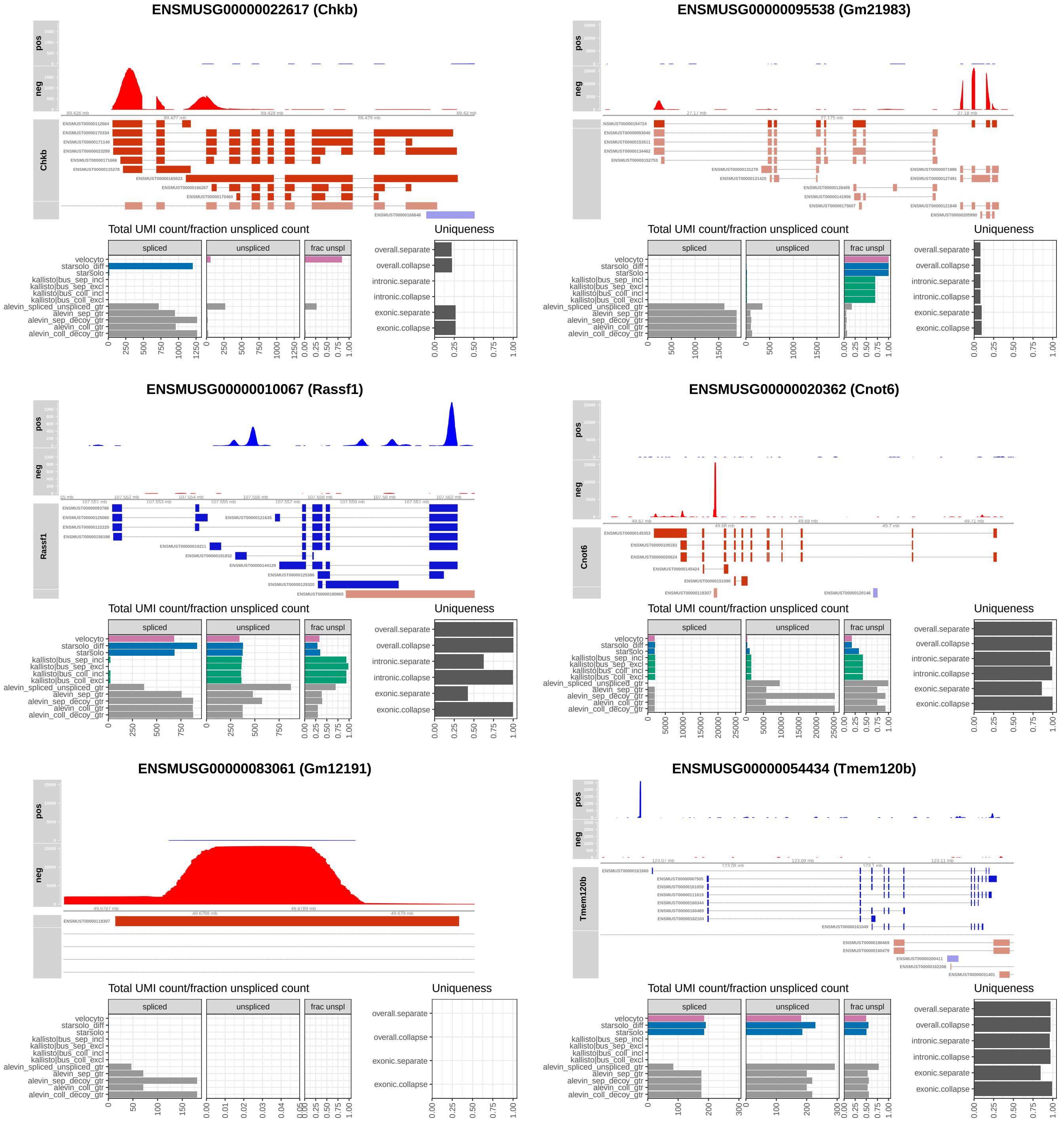


Figure S7: Examples of genes illustrating major differences among the quantification methods. The coverage tracks show the number of reads overlapping each base position, on the positive (blue) or negative (red) strand. The isoforms of the main gene in focus of each panel are similarly shown in blue (positive strand) or red (negative strand). Any overlapping features from other genes are shown in the bottom annotation track, colored in muted blue or red, depending on the annotated strand. The bottom panels show the total exonic and intronic UMI count assigned to the displayed gene by the different quantification methods, as well as the fraction of unspliced counts, and the fraction of unique k-mers in the gene overall, as well as in the exons and introns (see Methods).

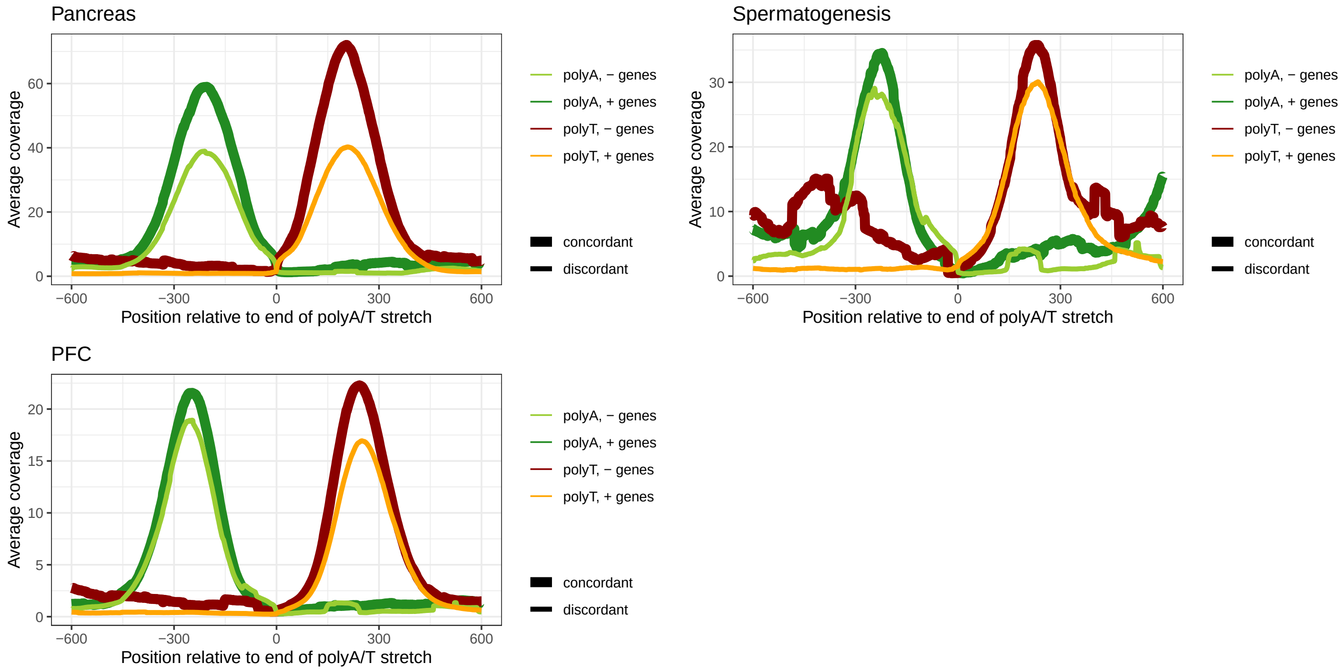


Figure S8: Average coverage of genomic regions around polyA/polyT stretches (at least 15 nt, with at most one mismatch per 15 nt) in introns of genes on the forward and reverse strands. Similarly to La Manno et al. (2018), we observe consistent coverage around discordant internal priming regions.

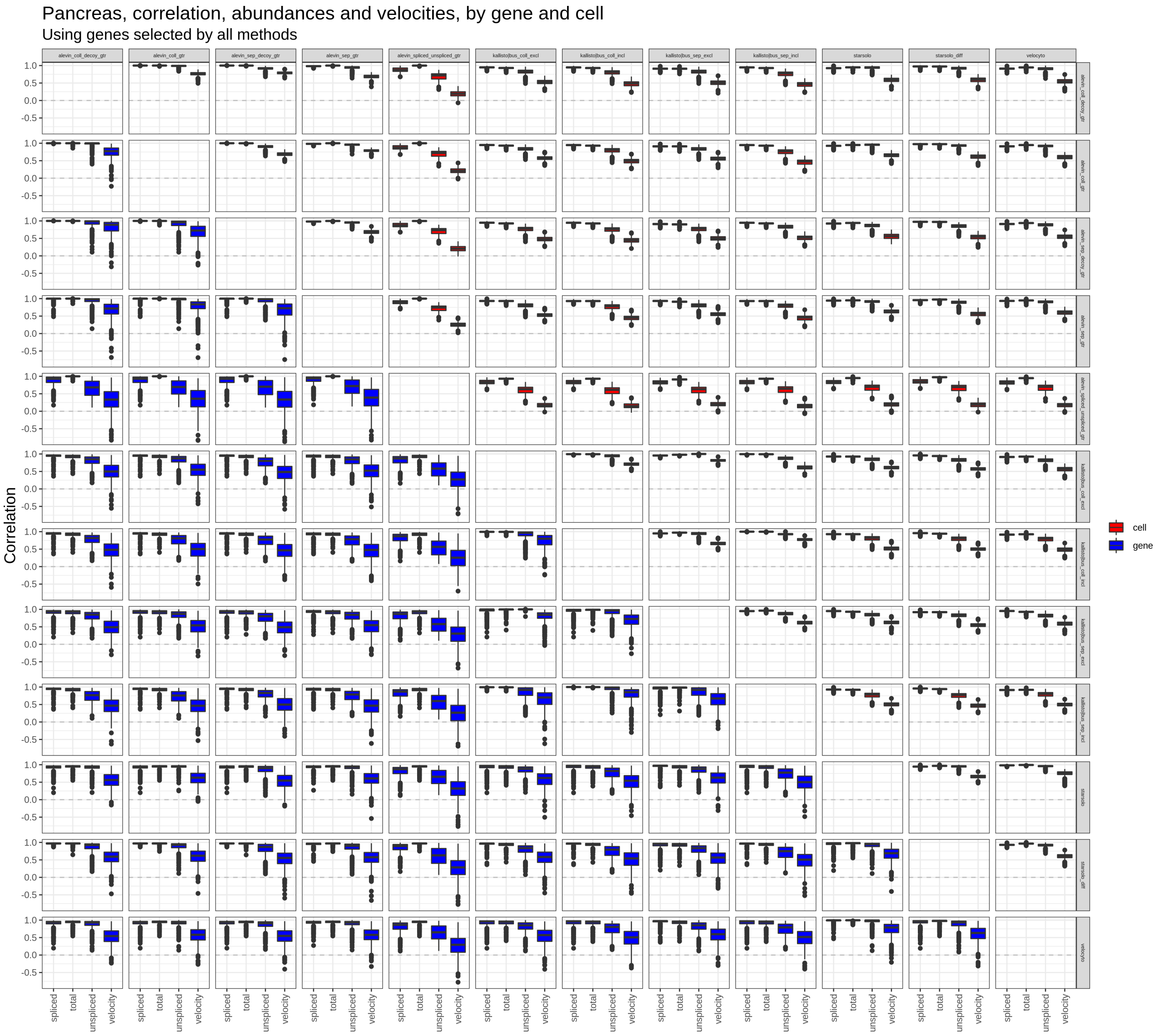


Figure S9: Cell- and gene-wise Spearman correlations of spliced, unspliced and total normalized abundance, as well as velocity estimates, between each pair of quantification methods.

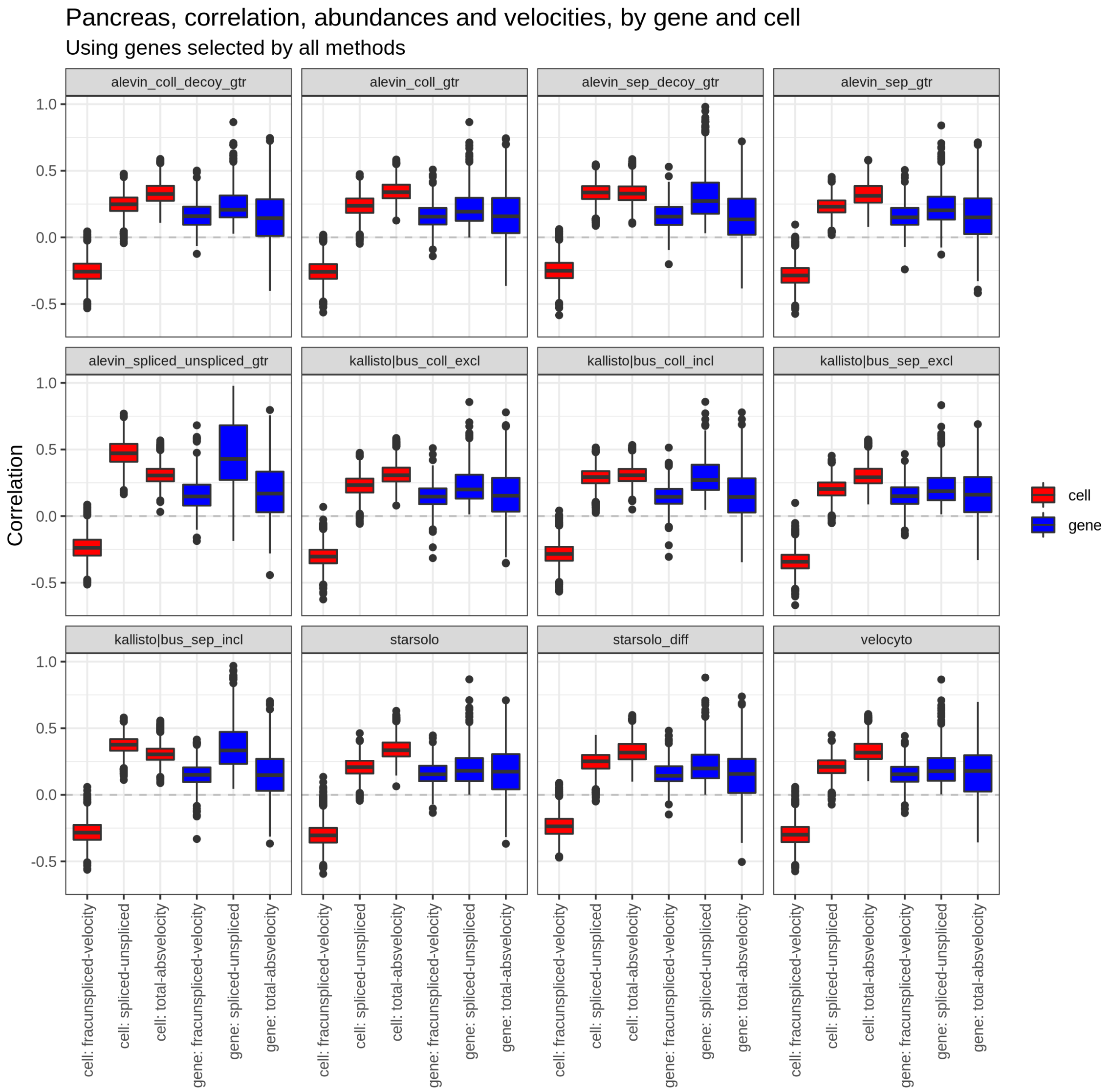


Figure S10: Spearman correlation between total normalized abundance (from *scVelo*) and estimated velocity, and between spliced and unspliced normalized abundances, either by gene or by cell, for all quantification methods.

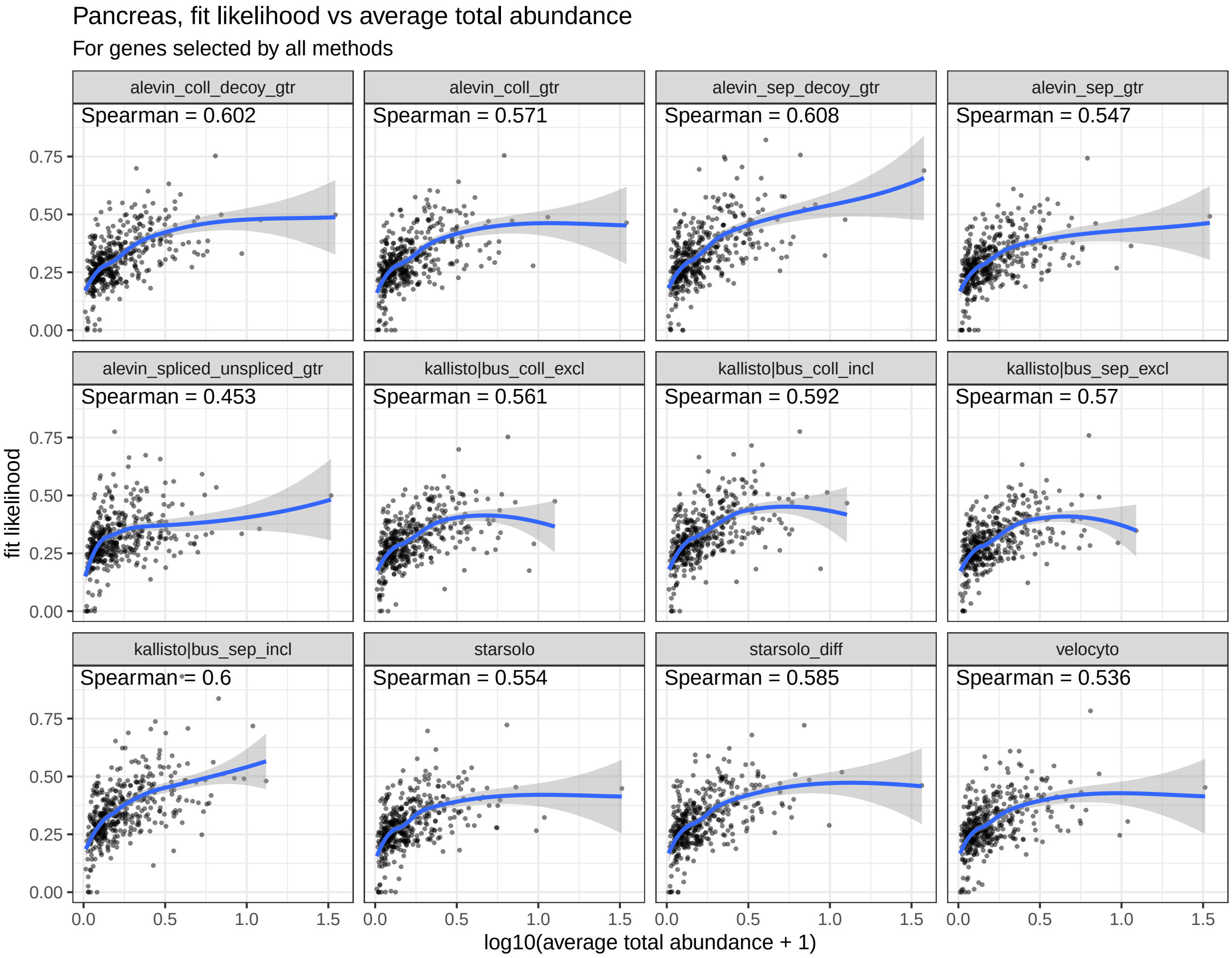


Figure S11: Spearman orrelation between average total normalized abundance across cells (from *scVelo*) and estimated likelihood of the velocity fit, for all quantification methods (Pancreas data).

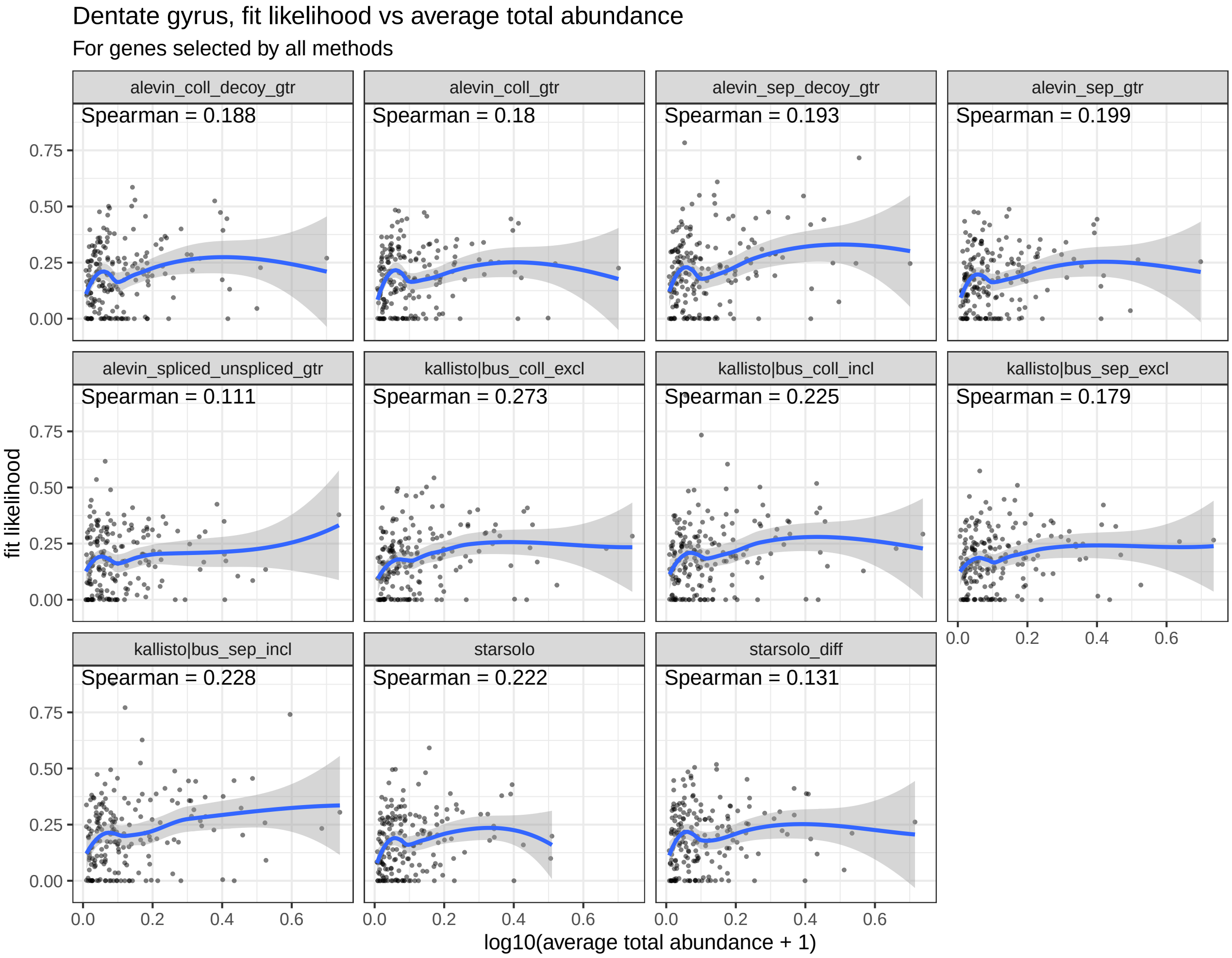


Figure S12: Spearman correlation between average total normalized abundance across cells (from *scVelo*) and estimated likelihood of the velocity fit, for all quantification methods (Dentate gyrus data).

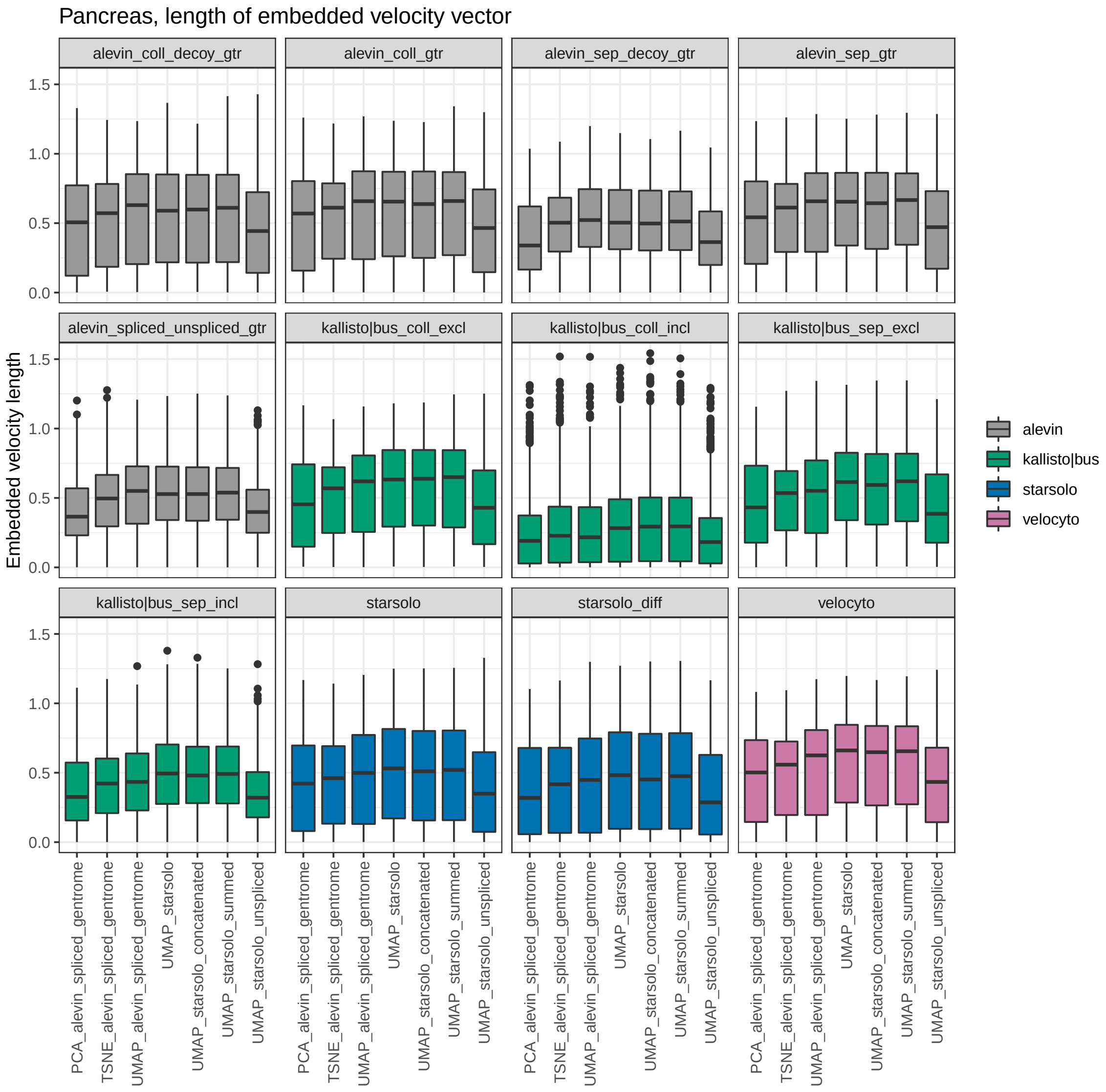


Figure S13: Distribution of the length of the embedded velocity vectors based on different quantification methods (panels), in different reduced dimension representations ($x$-axis), for the Pancreas data set. Since the displacement vectors in the low-dimensional representation are normalized before a weighted average is taken to determine the velocity embedding for a given cell, the length of the velocity embedding indicates to what extent the cells to which a cell $i$ has a high transition probability are all located in the same direction from cell $i$ in the low-dimensional representation. Consequently, the length of the embedded velocity vector provides one way of measuring how interpretable the low-dimensional representation is, from a velocity point of view. While overall only small differences are seen between dimension reductions, UMAP generally showed a small improvement in interpretability compared to PCA and TSNE, and using only the unspliced counts as the basis for the embedding was typically less informative.

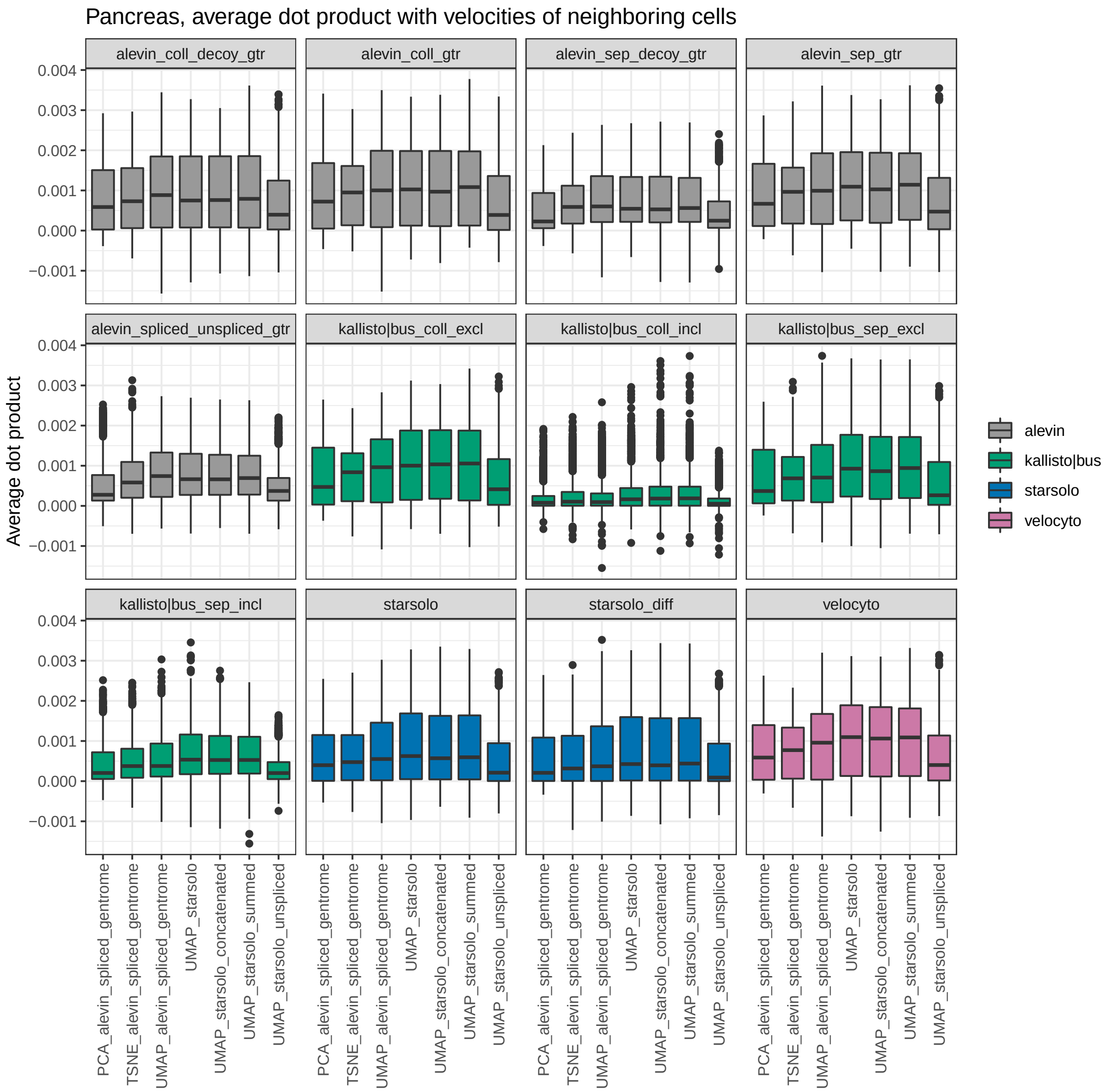


Figure S14: Distribution of the average similarity (dot product) between embedded velocity vectors among neighboring cells across a range of reduced dimension representations, for the Pancreas data set. A high similarity suggests that velocity streamlines in the reduced dimension representation are more easily interpretable, since they are designed to summarize the dynamics across neighbouring cells.

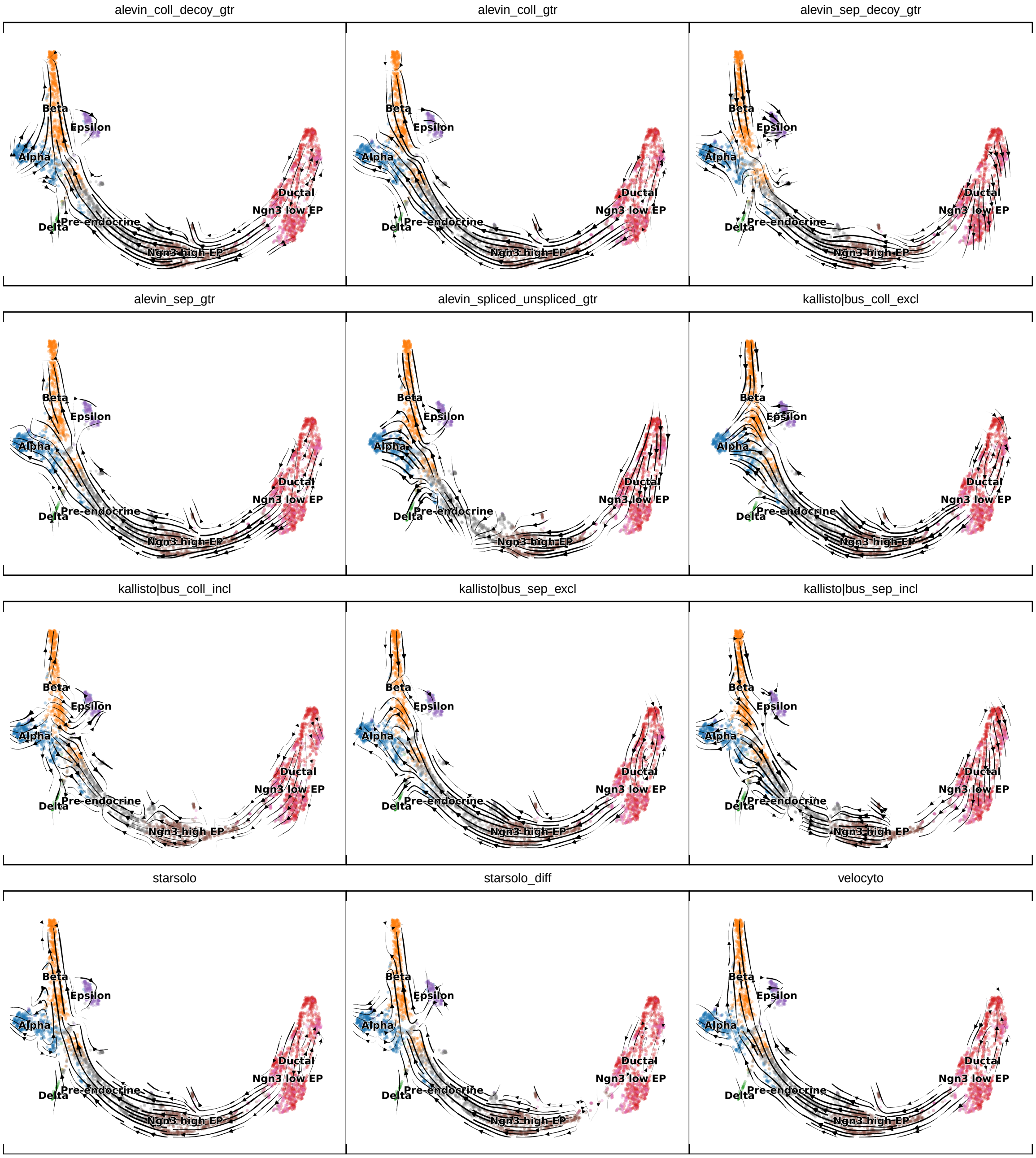


Figure S15: Inferred velocities in the pancreas data set, visualized on top of the same UMAP representation.

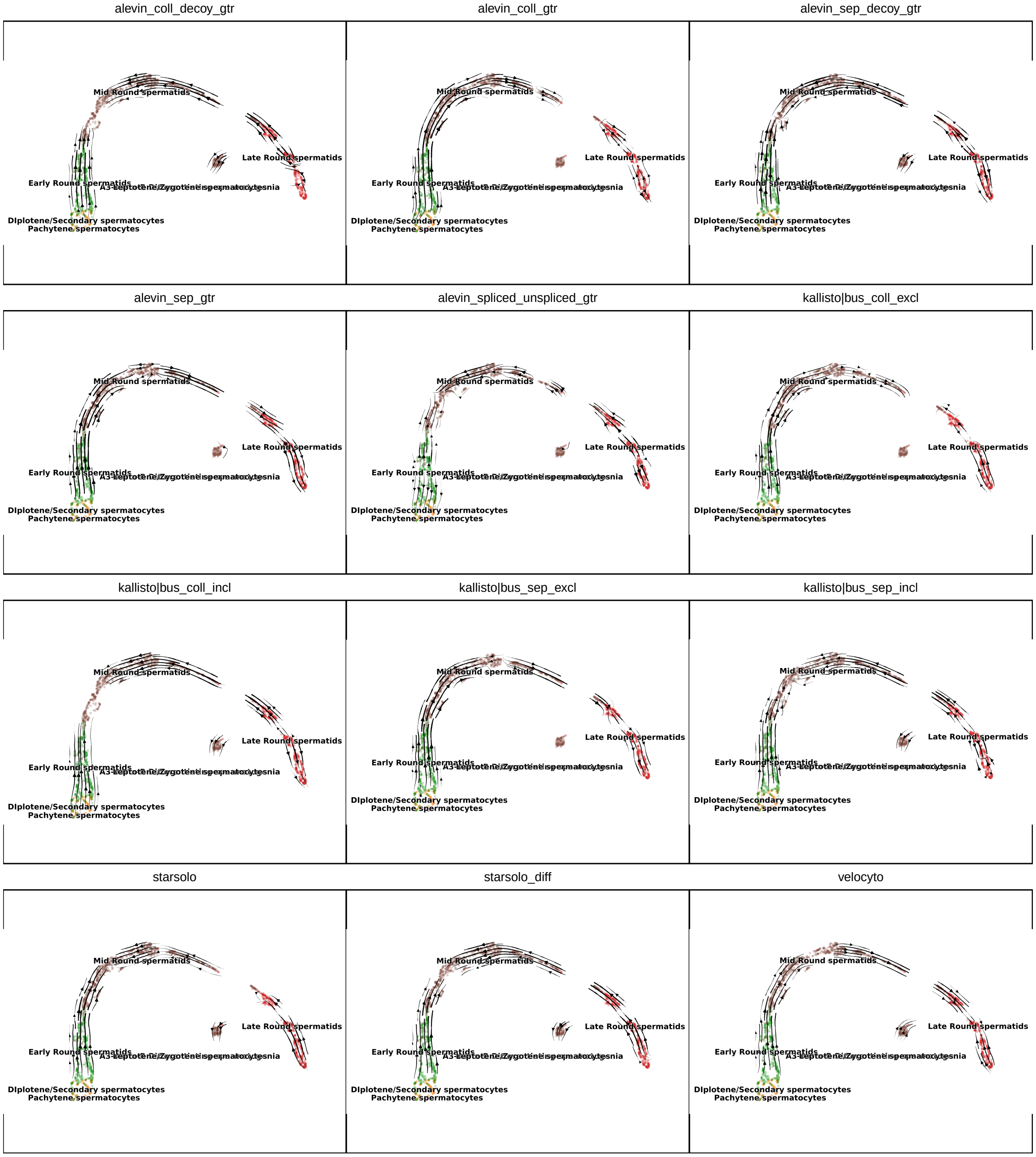


Figure S16: Inferred velocities in the spermatogenesis data set, visualized on top of the same UMAP representation.

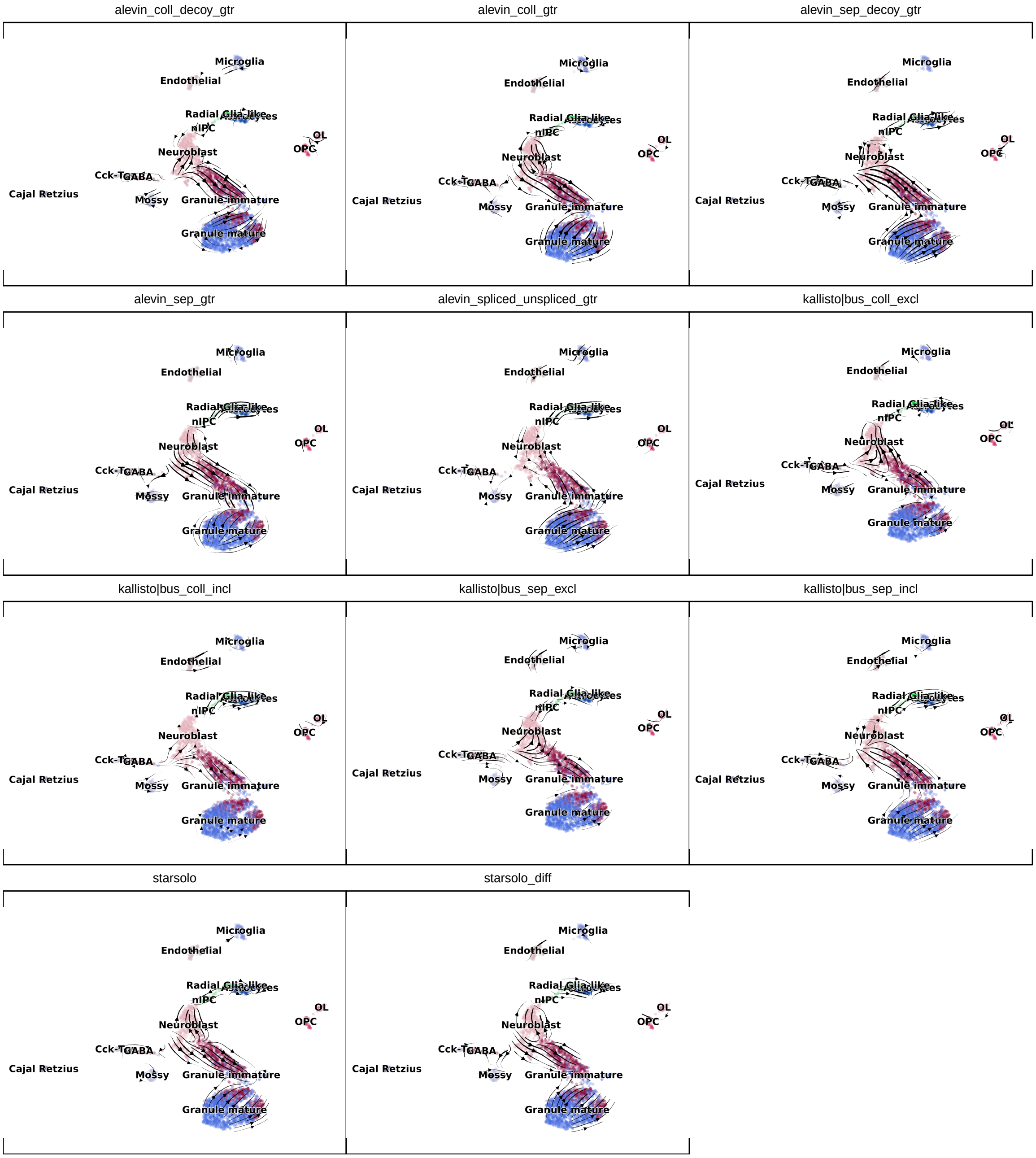


Figure S17: Inferred velocities in the dentate gyrus data set, visualized on top of the same UMAP representation.

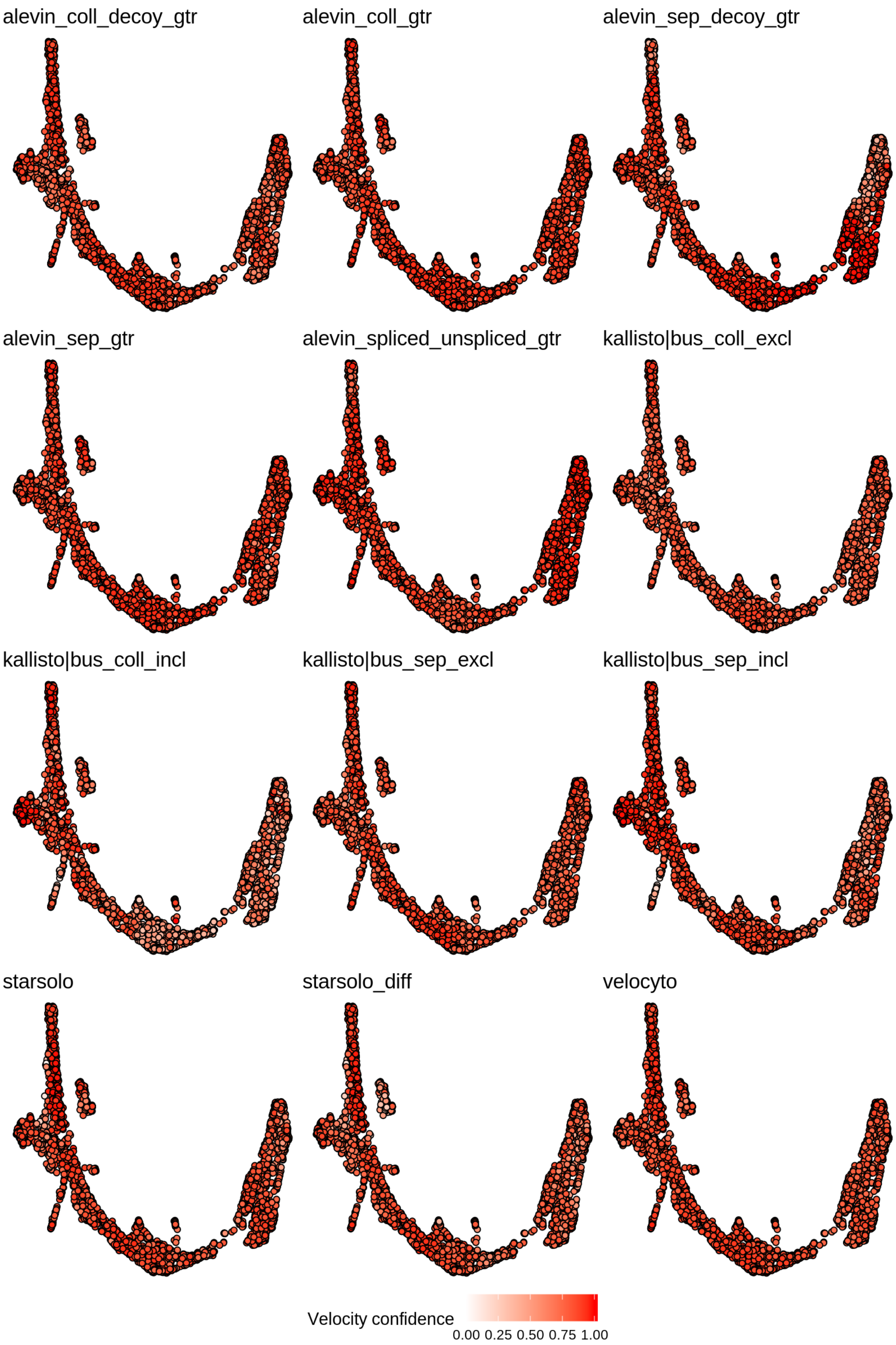


Figure S18: Velocity confidence estimates (similarity between the estimated velocities for a cell and those of its neighbors) obtained by *scVelo*, for the pancreas data set.

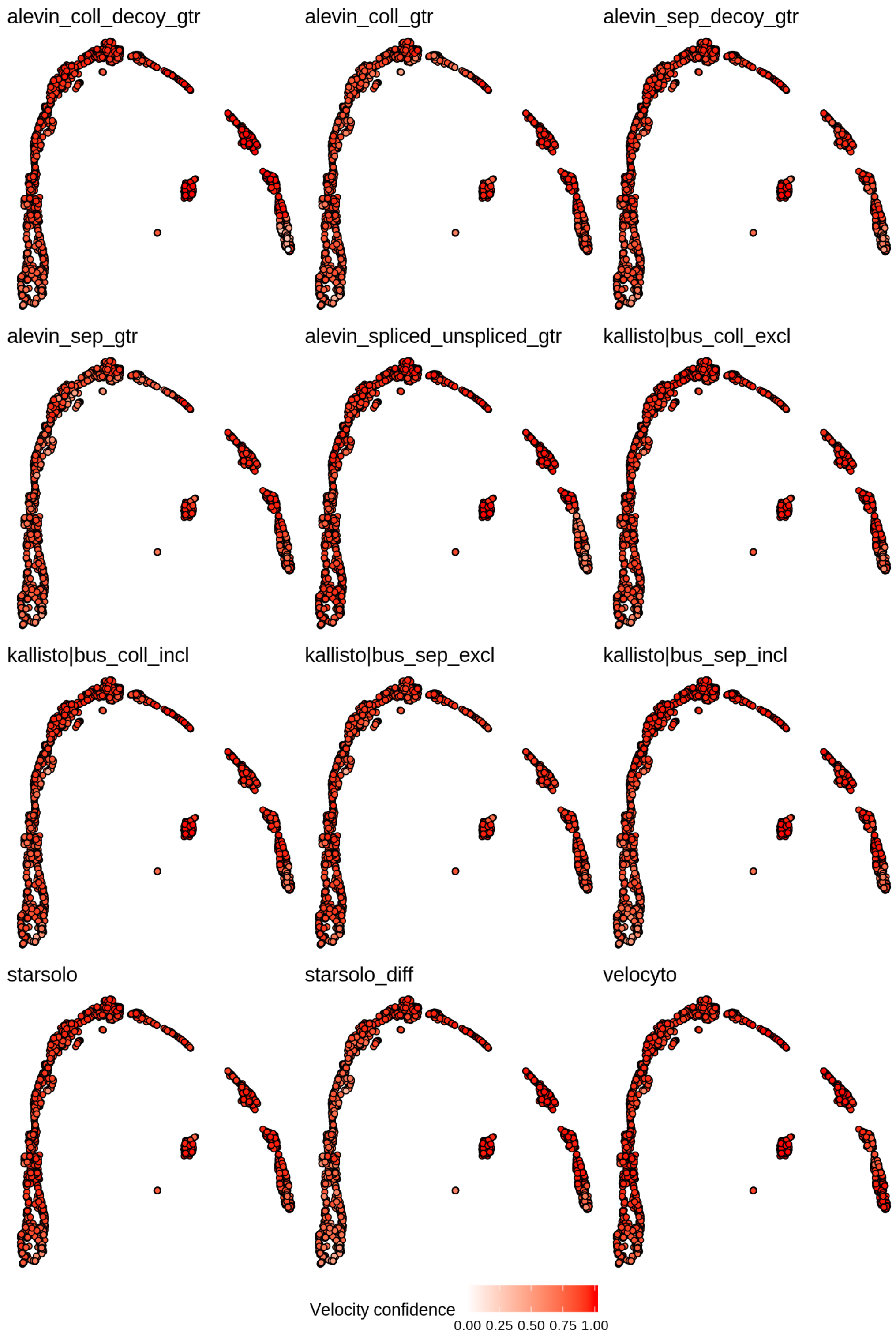


Figure S19: Velocity confidence estimates (similarity between the estimated velocities for a cell and those of its neighbors) obtained by *scVelo*, for the spermatogenesis data set.

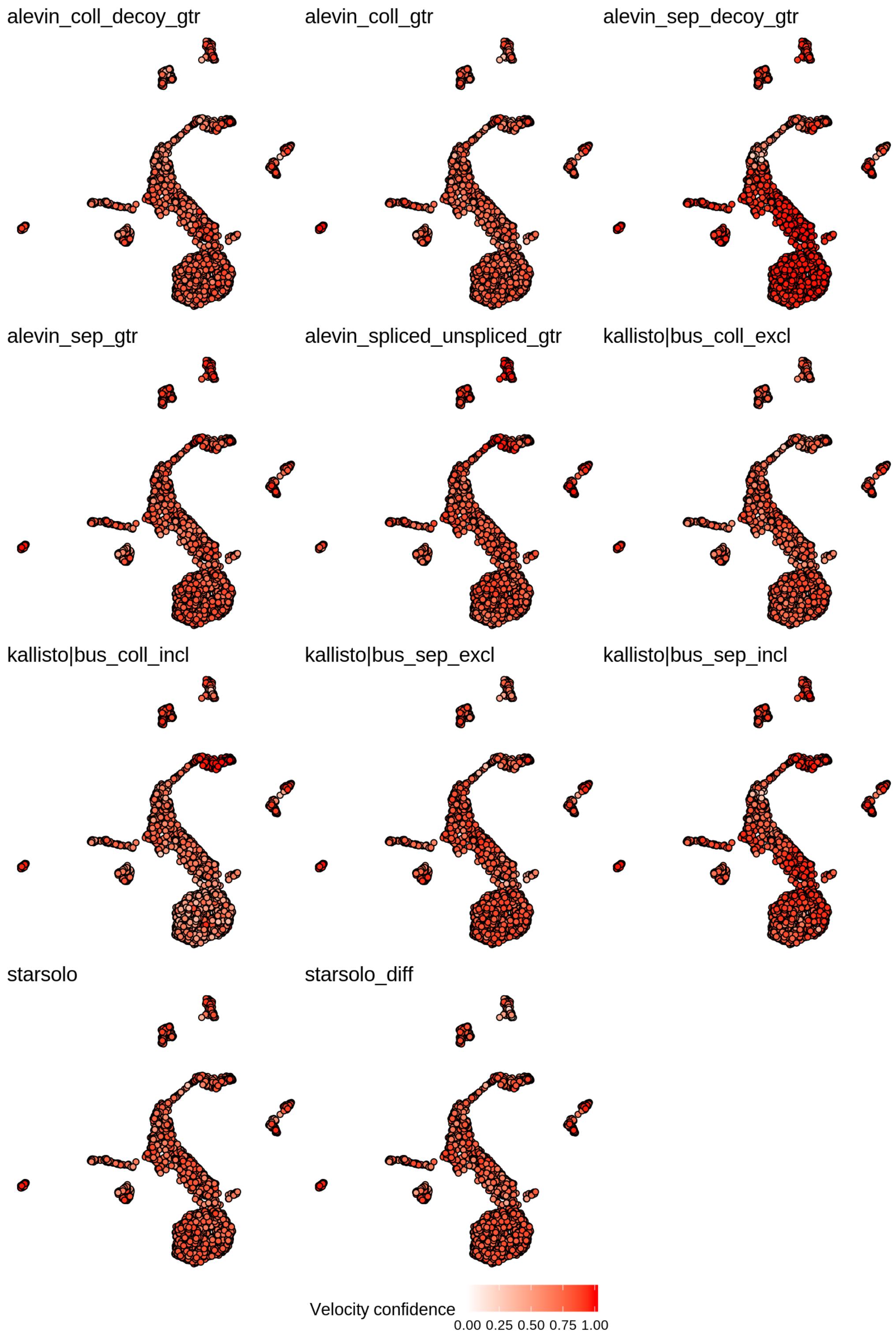


Figure S20: Velocity confidence estimates (similarity between the estimated velocities for a cell and those of its neighbors) obtained by *scVelo*, for the dentate gyrus data set.

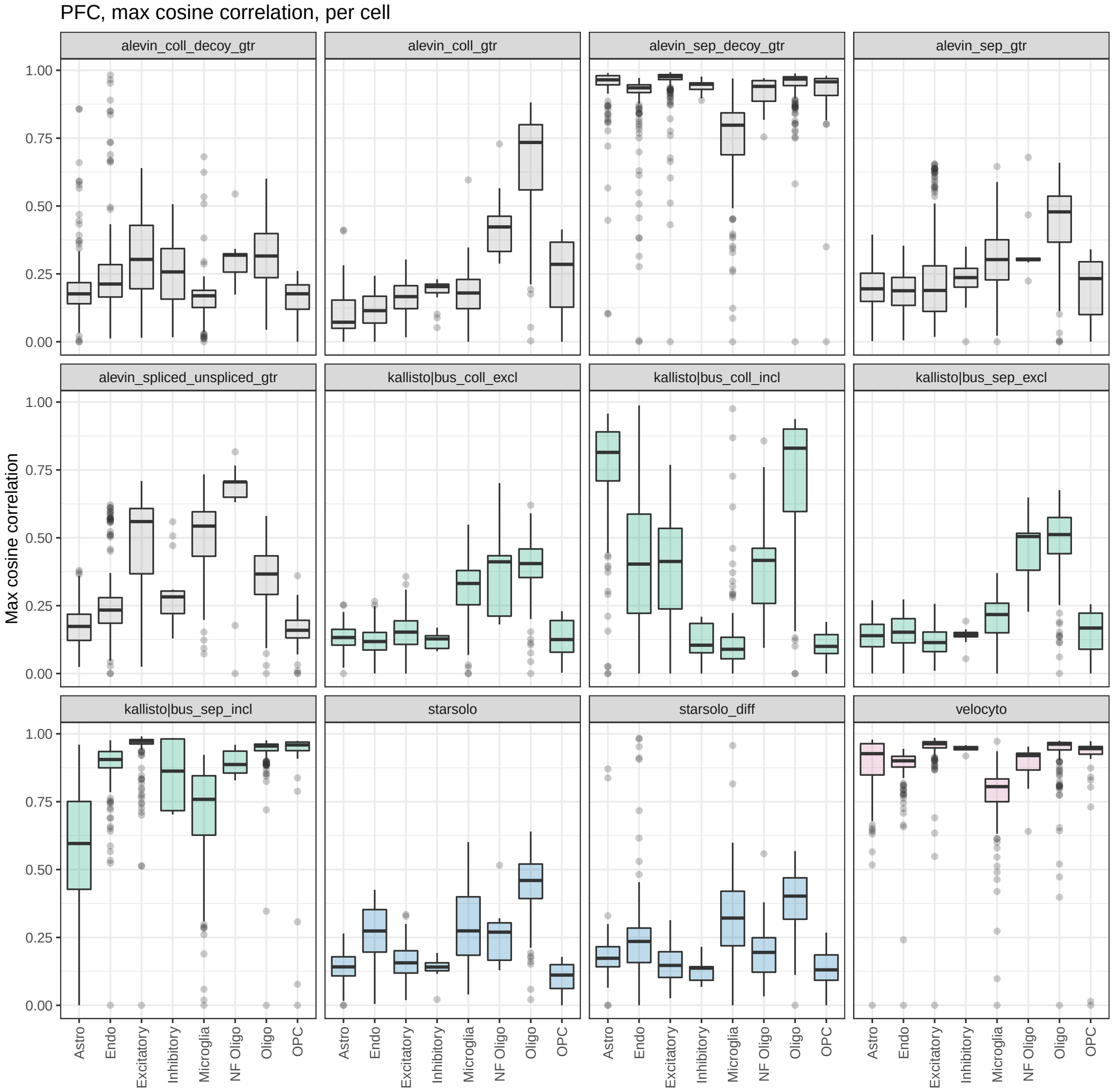


Figure S21: Maximal cosine correlation between the velocity vector and the displacement vector to other cells in the neighborhood, estimated by *scVelo* for different quantification methods in the PFC data set. A high cosine correlation for a cell *i* indicates that the estimated velocity for the cell points in the direction of some other cell in the data set.

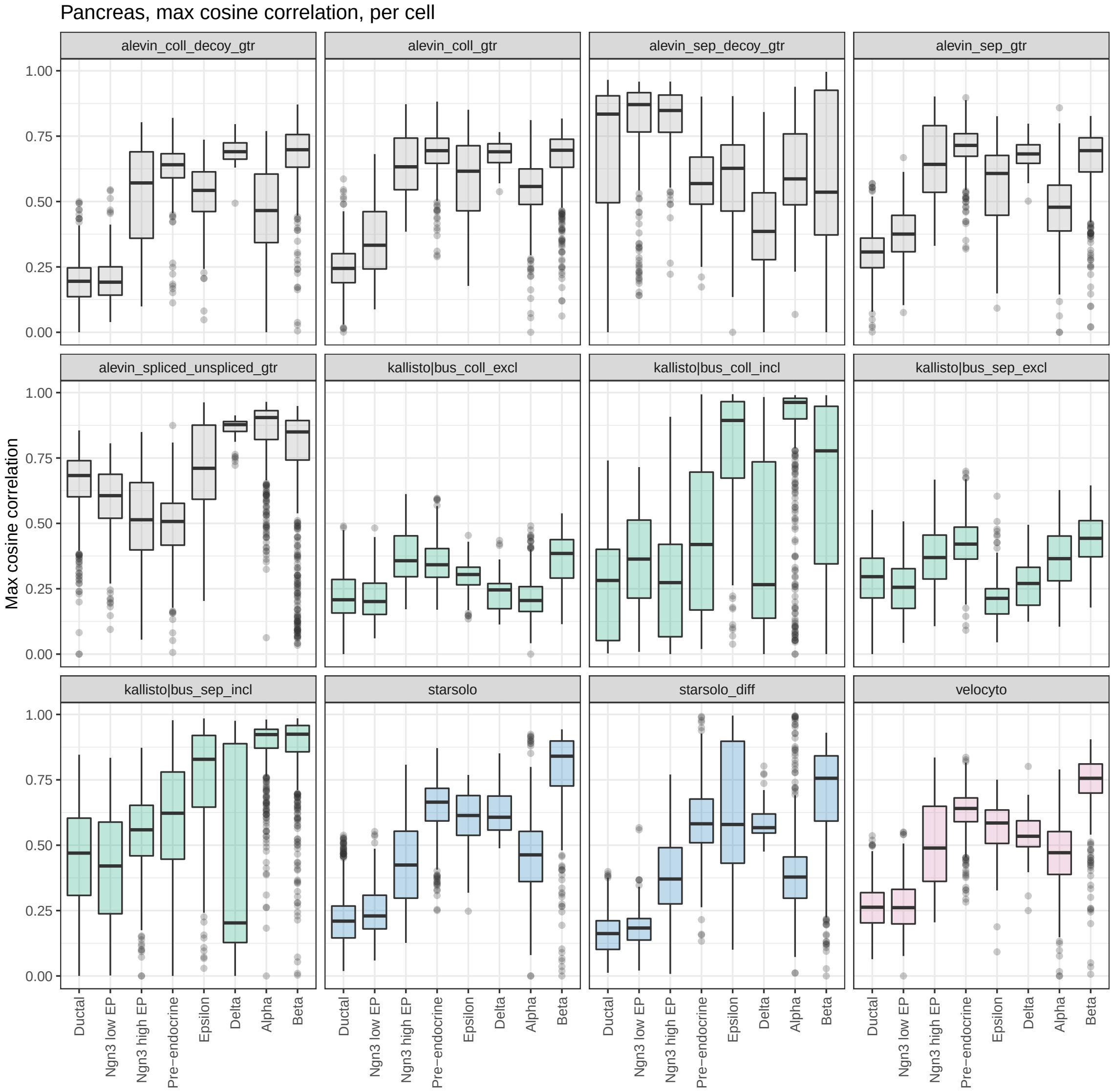


Figure S22: Maximal cosine correlation between the velocity vector and the displacement vector to other cells in the neighborhood, estimated by *scVelo* for different quantification methods in the pancreas data set. A high cosine correlation for a cell *i* indicates that the estimated velocity for the cell points in the direction of some other cell in the data set.

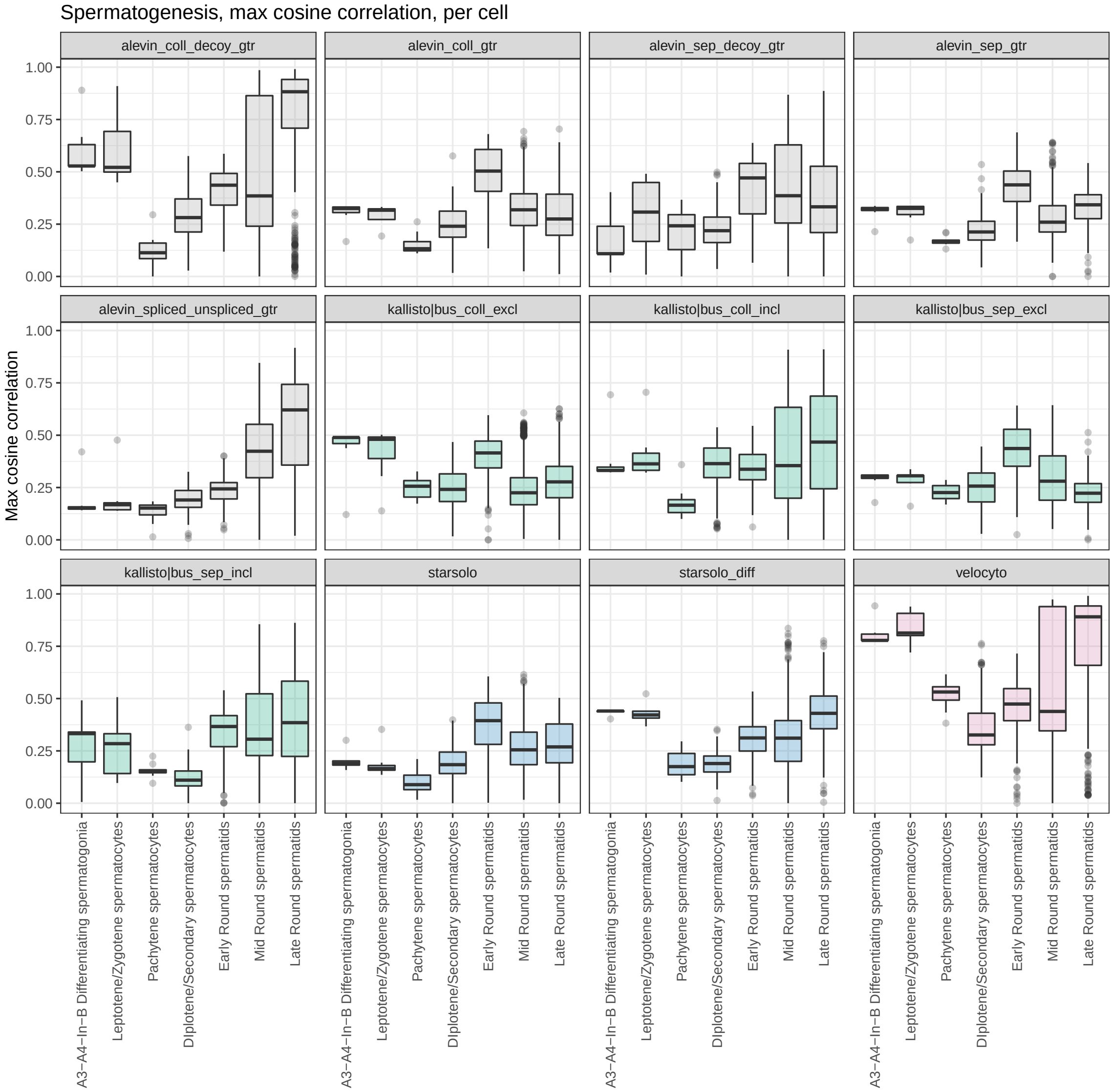


Figure S23: Maximal cosine correlation between the velocity vector and the displacement vector to other cells in the neighborhood, estimated by *scVelo* for different quantification methods in the spermatogenesis data set. A high cosine correlation for a cell $i$ indicates that the estimated velocity for the cell points in the direction of some other cell in the data set.

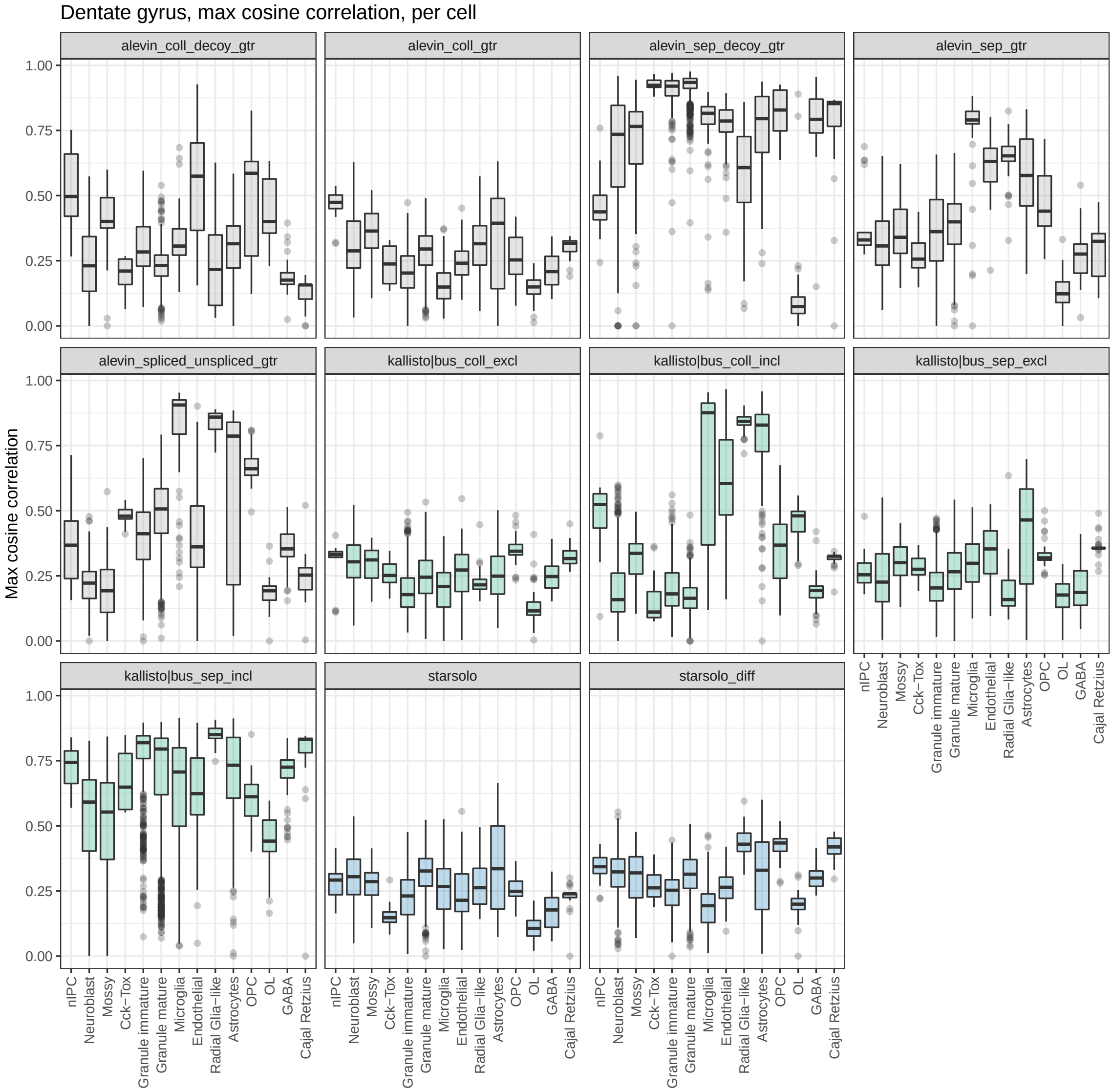


Figure S24: Maximal cosine correlation between the velocity vector and the displacement vector to other cells in the neighborhood, estimated by *scVelo* for different quantification methods in the dentate gyrus data set. A high cosine correlation for a cell *i* indicates that the estimated velocity for the cell points in the direction of some other cell in the data set.

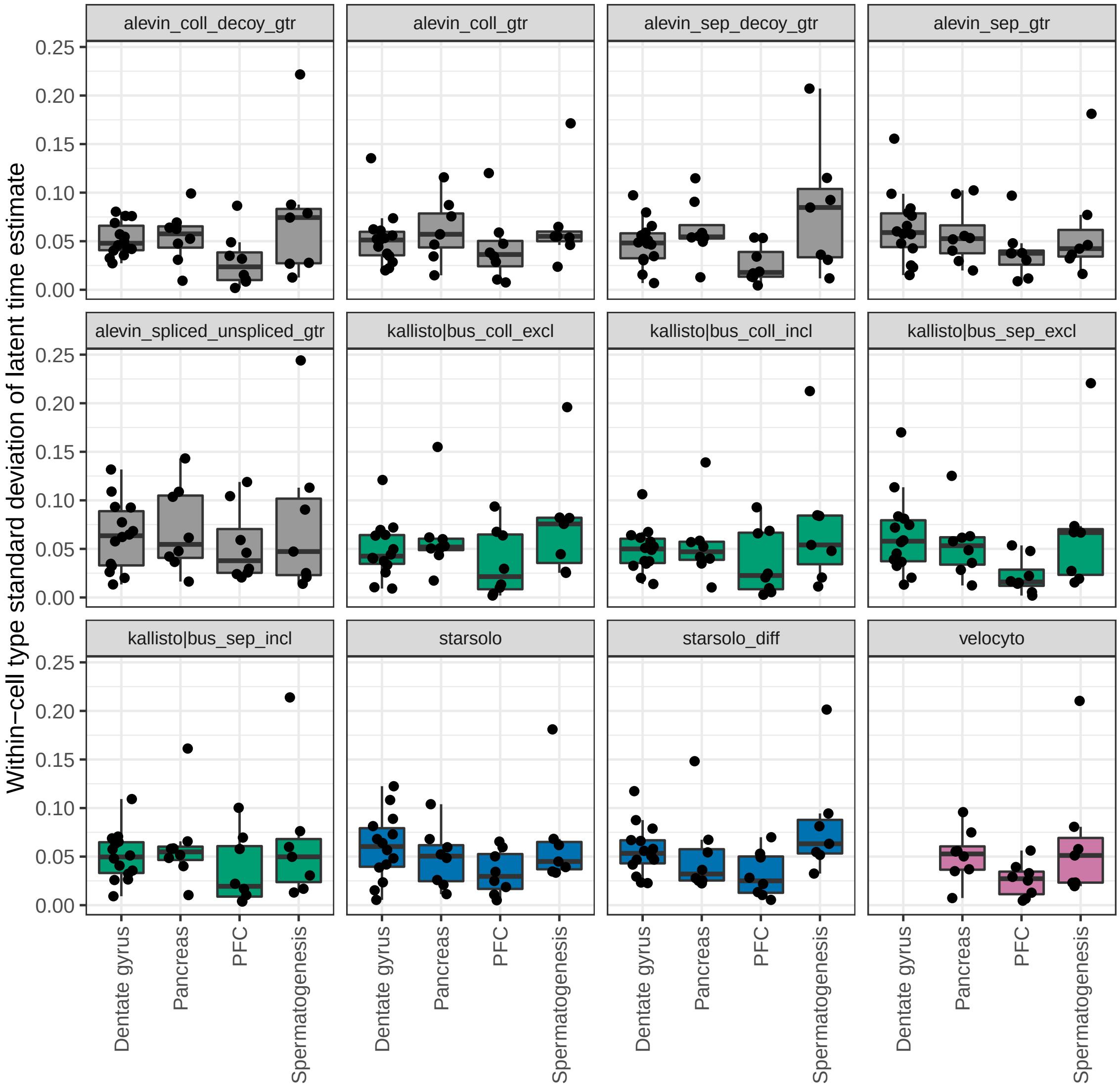


Figure S25: Standard deviation of estimated latent time within each cell type, across the four data sets. Each dot represents one cell type. This is used as a proxy for the amount of continuous dynamics in the data set, and expected to be lower for the PFC data set than for the others.
